# The Hippo kinases control inflammatory Hippo signaling and restrict bacterial infection in eukaryotic phagocytes

**DOI:** 10.1101/2023.12.08.570858

**Authors:** Brendyn M. St. Louis, Sydney M. Quagliato, Yu-Ting Su, Gregory Dyson, Pei-Chung Lee

## Abstract

The Hippo kinases MST1 and MST2 initiate a highly conserved signaling cascade called the Hippo pathway that limits organ size and tumor formation in animals. Intriguingly, pathogens hijack this host pathway during infection, but the role of MST1/2 in innate immune cells against pathogens is unclear. In this study, we generated *Mst1/2* knockout macrophages to investigate the regulatory activities of the Hippo kinases in immunity. Transcriptomic analyses identified differentially expressed genes (DEGs) that are enriched in biological pathways, such as systemic lupus erythematosus, tuberculosis, and apoptosis. Surprisingly, pharmacological inhibition of the downstream components LATS1/2 in the canonical Hippo pathway did not affect expression of a set of immune DEGs, suggesting that MST1/2 control these genes via alternative inflammatory Hippo signaling. Moreover, MST1/2 may affect immune communication by influencing the release of cytokines, such as TNFα, CXCL10, and IL-1ra. Comparative analyses of the single- and double-knockout macrophages revealed that MST1 and MST2 differentially regulate TNFα release and expression of the immune transcription factor, MAF, demonstrating that the two homologous Hippo kinases individually play a unique role in innate immunity. Notably, MST1 and MST2 are both required for macrophages to activate apoptosis. Lastly, we demonstrated that the Hippo kinases are critical factors in mammalian macrophages and single-cell amoebae to restrict infection by *Legionella pneumophila*, *Escherichia coli*, and *Pseudomonas aeruginosa*. Together, these results uncover non-canonical inflammatory Hippo signaling in macrophages and the evolutionarily conserved role of the Hippo kinases in anti-microbial defense of eukaryotic hosts.

## Introduction

The Hippo pathway exists in all eukaryotes, including single-cell amoebae and multicellular organisms like humans (1), and controls important biological functions, such as cell cycle progression and organ development (2, 3). In canonical Hippo signaling of mice and humans, mammalian STE20-like protein kinases-1 and -2 (MST1 and MST2) are the Hippo kinases that phosphorylate the scaffold protein, Mps one binder kinase activator-like 1A and B (MOB1A/B), and the downstream kinases, large tumor suppressor homolog 1 and 2 (LATS1/2). Phosphorylated MOB1 and LATS1/2 form active kinase complexes which subsequently phosphorylate the transcription regulators, Yes-associated protein 1 (YAP1) and WW domain-containing transcription regulator protein 1 (WWTR1). Once phosphorylated, YAP1 and WWTR1 are sequestered in the cytoplasm or undergo proteasomal degradation. Conversely, when canonical Hippo signaling is off, unphosphorylated YAP1/WWTR1 shuttle into the nucleus and interact with the transcription enhancer factors TEADs to control gene expression (2–6).

While *Mst1/2* double knockout (*Mst1/2^−/−^*) leads to embryonic lethality (7), conditional *Mst1/2^−/−^* in the liver or intestines causes enlarged tissues and tumor formation in mouse models (8–11). Ectopic overproduction of MST1 or MST2 in cell lines promotes apoptosis (12–16), and elevated YAP1 protein levels are associated with human cancers (17), suggesting that MST1/2 possess pro-apoptotic activities to limit cell proliferation and tumors. However, naïve T cells isolated from *Mst1/2^−/−^* mice and humans with defective MST1 are highly sensitive to apoptosis (18–20), indicating that the regulatory activities of MST1/2 in programmed cell death depend on specific cell types and conditions. While being one of the key pathways in human cancer research (21), the Hippo pathway also has an emerging role in immunity. Humans with loss-of-function mutations in the *MST1* gene have clinical histories of recurrent infections and pneumonia (19, 20, 22). Mice with conditional *Mst1/2^−/−^* in myeloid cells are more susceptible to death caused by septic peritonitis (23). However, it remains undefined if MST1 and MST2 have shared or distinct roles in host defense. Meanwhile, the importance of the Hippo pathway in microbial pathogenesis is rising. The opportunistic pathogen *Legionella pneumophila* uses LegK7, a bacterial effector protein, to hijack the Hippo scaffold protein MOB1 (24). Similarly, *Chlamydia trachomatis* and *Salmonella enterica* use the effector proteins Tarp and SopB, respectively, to manipulate the Hippo transcription regulator YAP1 (25, 26). Moreover, human papilloma virus E6 protein targets LATS kinases and TEAD transcription factor to promote pathogenesis (27, 28), and replication of the SARS-CoV2 virus is increased in host cells with MST1/2 knockdown (29).

Here, we use comparative genetic approaches to investigate the roles of MST1/2 in macrophages, the professional phagocytes in innate immunity. We found that MST1/2 are critical regulators for expression of hundreds of macrophage genes involved in immune disorders and cell death pathways. Interestingly, MST1/2 regulate a set of immune genes through non-canonical Hippo signaling cascades that are independent of the canonical Hippo pathway. MST1/2 have different regulatory effects on the release of cytokines in macrophages, indicating that the Hippo kinases affect cell-cell communication in immunity. Lastly, we demonstrate the requirement of MST1/2 in programmed cell death of macrophages and characterize the conserved Hippo kinases as critical host factors in macrophages and protozoans to restrict invading bacterial pathogens. These findings uncover an inflammatory Hippo signaling pathway in innate immunity that is unique and distinct from the canonical Hippo pathway in organ development.

## Results

### The macrophage transcriptome controlled by MST1/2

To investigate the role of the Hippo kinases in immune cells, we knocked out both *Mst1* and *Mst2* genes (*Mst1/2^−/−^*) in mouse immortalized bone marrow-derived macrophages (iBMDMs) using CRISPR-Cas9 genome-editing techniques. We obtained multiple *Mst1/2^−/−^* iBMDM clones (N5, N12, N13, and N19) that had no detectable levels of MST1 and MST2 (Fig. 1A). In these clones, the protein levels of LATS1 and MOB1 remained comparable to the levels in wildtype (WT) iBMDMs, but MOB1 phosphorylation was abolished since MOB1 is a known substrate of MST1/2, further confirming that MST1/2 were absent in these cells (Fig. 1A). Next, we used RNA sequencing (RNAseq) to profile global gene expression in iBMDMs. WT and *Mst1/2^−/−^* iBMDMs were either treated with or without lipopolysaccharide (LPS), the outer membrane component of Gram-negative bacteria, to evaluate the effects of MST1/2 on macrophage gene expression under non-stimulated and stimulated conditions. The biological coefficient of variation plot showed that all four *Mst1/2^−/−^* clones clustered relatively close together and were markedly distinct from the WT macrophages (Fig. 1B). LPS stimulation caused similar shifts in WT and *Mst1/2^−/−^* macrophages (Fig. 1B). This global analysis showed that the *Mst1/2^−/−^* clones behave similarly, and MST1/2 have significant impacts on macrophage gene expression.

**Figure 1.**
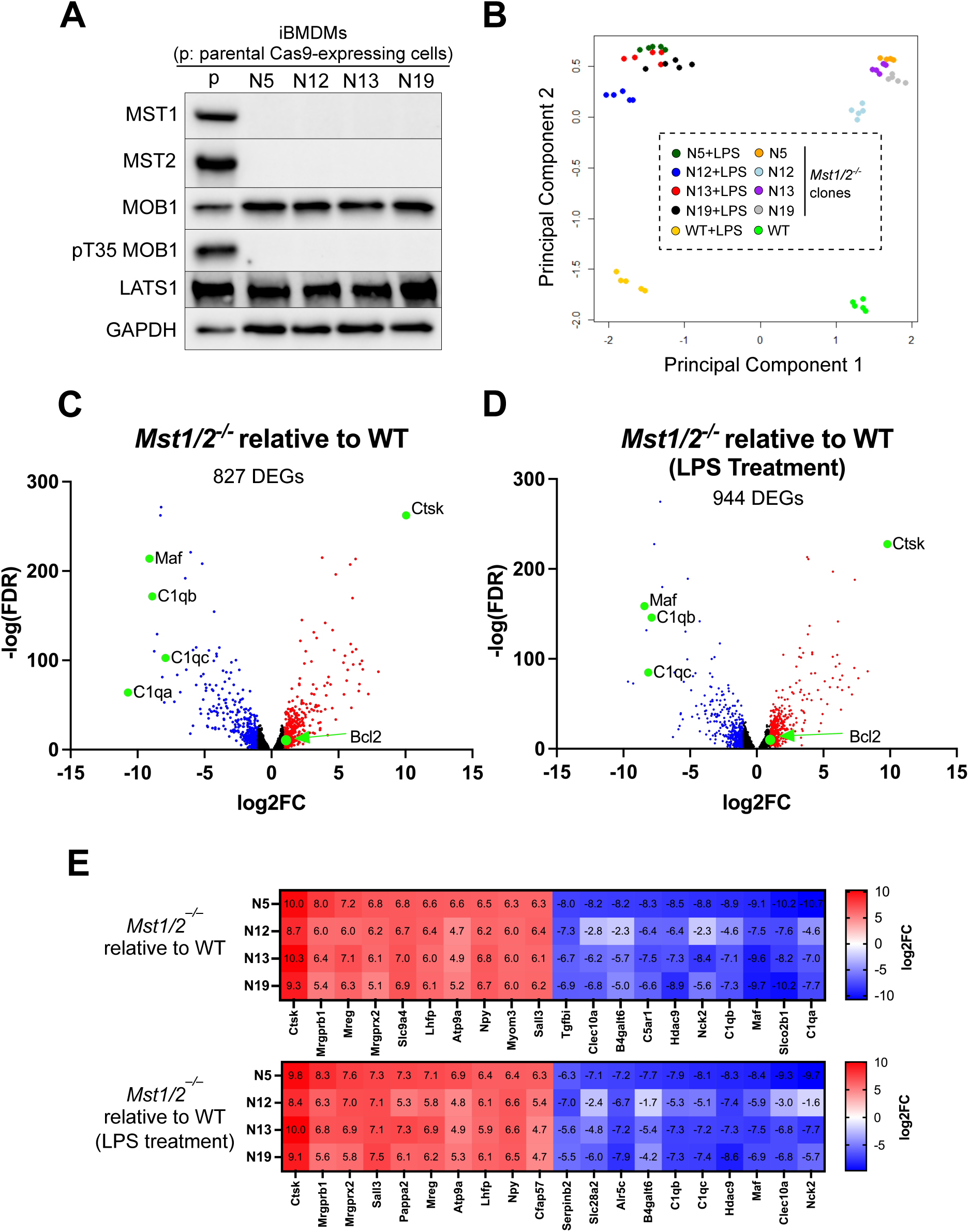
Global gene transcription controlled by MST1/2 in macrophages. **A**. Loss of MST1/2 proteins and MOB1 phosphorylation in *Mst1/2^−/−^* iBMDMs. Cell lysate from the iBMDMs clones were analyzed by immunoblotting for the indicated proteins and phosphorylation of the threonine 35 residue on MOB1 (pT35 MOB1). GADPH was used as an internal control. **B**. Principal component analysis of gene expression profile similarities between wildtype (WT) iBMDMs and *Mst1/2^−/−^* clones treated with or without 2 µg/mL of LPS for three hours. Data represent samples collected from five independent repeats. **C and D**. Volcano plots for gene expression of the representative *Mst1/2^−/−^* clone N5 compared to WT iBMDMs based on RNAseq data. Differentially expressed genes (DEGs) were selected with greater than 100 transcript reads in either WT or *Mst1/2^−/−^* N5, log2-fold change (log2FC) greater than ± 1 and false discovery rate (FDR) <0.01. Red, up-regulated DEGs; blue, down-regulated DEGs. Green spots labeled the genes further analyzed or discussed in this study. **E**. Expression heat map of the top 10 up- or down-DEGs in all four *Mst1/2^−/−^* clones. The numbers within the color boxes indicate the log2-fold changes vs. WT iBMDMs.

Comparing the *Mst1/2^−/−^* N5 clone with the WT iBMDMs using these criteria; average reads>100 in N5 or WT; log2 fold-change >1 or <-1; false discover rate <0.01, 827 genes in no LPS condition and 944 genes in the LPS stimulation condition were identified as differentially expressed genes (DEGs) (Fig. 1C and 1D. Supplementary Table 1). Most of the top 10 up- or down-regulated genes had greater than 64-fold changes (log2-fold change > 6 or <-6), suggesting that MST1/2 are critical regulators for these genes (Fig. 1E). Many of the top DEGs were shared between the with and without LPS conditions (Fig. 1C-E). Interestingly, none of the top DEGs were characterized as the signature genes regulated by YAP1 and WWTR1, the downstream transcription regulators in the canonical Hippo pathway (30). We further examined the expression levels of twenty-two YAP1/WWTR1 signature genes identified in cancers by the previous study (30). Surprisingly, majority of the signature genes (17 out of 22) were poorly expressed in iBMDMs (Fig. S1A), including *Ctgf* and *Cyr61* which are well-known marker genes controlled by YAP1/WWTR1. Among the five genes that had relatively high expression levels (average reads>100), only two genes, *Dock5* and *Myof*, were moderately up-regulated (log2 fold-change: 1.12∼1.7) (Fig. S1B). Therefore, many genes in macrophages are possibly controlled through a non-canonical signaling cascade.

### Pathways, biological processes, and molecular signatures in macrophages controlled by MST1/2

To characterize the individual and combined interaction effects of *Mst1/2* gene knockouts and LPS stimulation, the gene lists were condensed and selected by more stringent criteria, (see Materials and Methods), resulting in 360 significant genes for *Mst1/2^−/−^*vs. WT iBMDMs, 706 significant genes for LPS vs. no LPS treatments, and 74 significant genes for combined interaction analysis. The significant genes were submitted to iPathwayGuide (Advaita Bioinformatics) for enrichment analyses, including pathways, gene ontology and biological processes. The iPathwayGuide confirmed that these macrophages responded to LPS stimulation since LPS was identified as the most significant upstream chemical from the LPS vs. no LPS significant genes (Fig. S2A). Marker genes of LPS stimulation, such as inducible NO synthase (*Nos2*), C-C motif chemokine ligand 5 (*Ccl5*) and interleukin 1α (*Il1a*), were highly induced in WT macrophages and all *Mst1/2^−/−^*clones (Fig. S2B), suggesting that *Mst1/2^−/−^* macrophages are responsive to LPS.

Next, we performed Gene Ontology (GO) analysis of the 360 significant genes from *Mst1/2^−/−^* vs. WT iBMDMs. While regulation of epithelial cell migration was the most significant GO biological process, several processes involved in programmed cell death, such as apoptosis, were also among the top biological processes (Fig. 2A). Regulation of cytokine production (GO:0001817) was also significantly enriched (p value=0.022), implying that MST1/2 might affect this process (Fig. 2A). Expression heat maps showed that the changes of genes involved in these processes were similar among the *Mst1/2^−/−^* clones (Fig. 2B). Analyzing the combined *Mst1/2* knockout and LPS interaction effect identified immune responses, cytokine production and programmed cell death as the top GO biological processes, with most of the processes being involved in immune activities (Fig. 2C). Likewise, the pathway analyses revealed that the genes from the *Mst1/2^−/−^* vs. WT comparison were highly enriched in immune pathways, such as tuberculosis, systemic lupus erythematosus, and apoptosis (Fig. 2D). Notably, the identified pathways included the FoxO signaling pathway which has been shown in other cell types to be influenced by MST1/2 (31–33) (Fig. 2D). Consistent with the GO analyses, the TNF signaling pathway and cytokine-cytokine receptor interaction were the two most enriched pathways in the interaction effects of *Mst1/2* knockout and LPS treatment (Fig. 2E). Together, these bioinformatic results illustrated a role for MST1/2 in the regulation of gene expression related to a variety of inflammatory responses and programmed cell death in macrophages.

**Figure 2.**
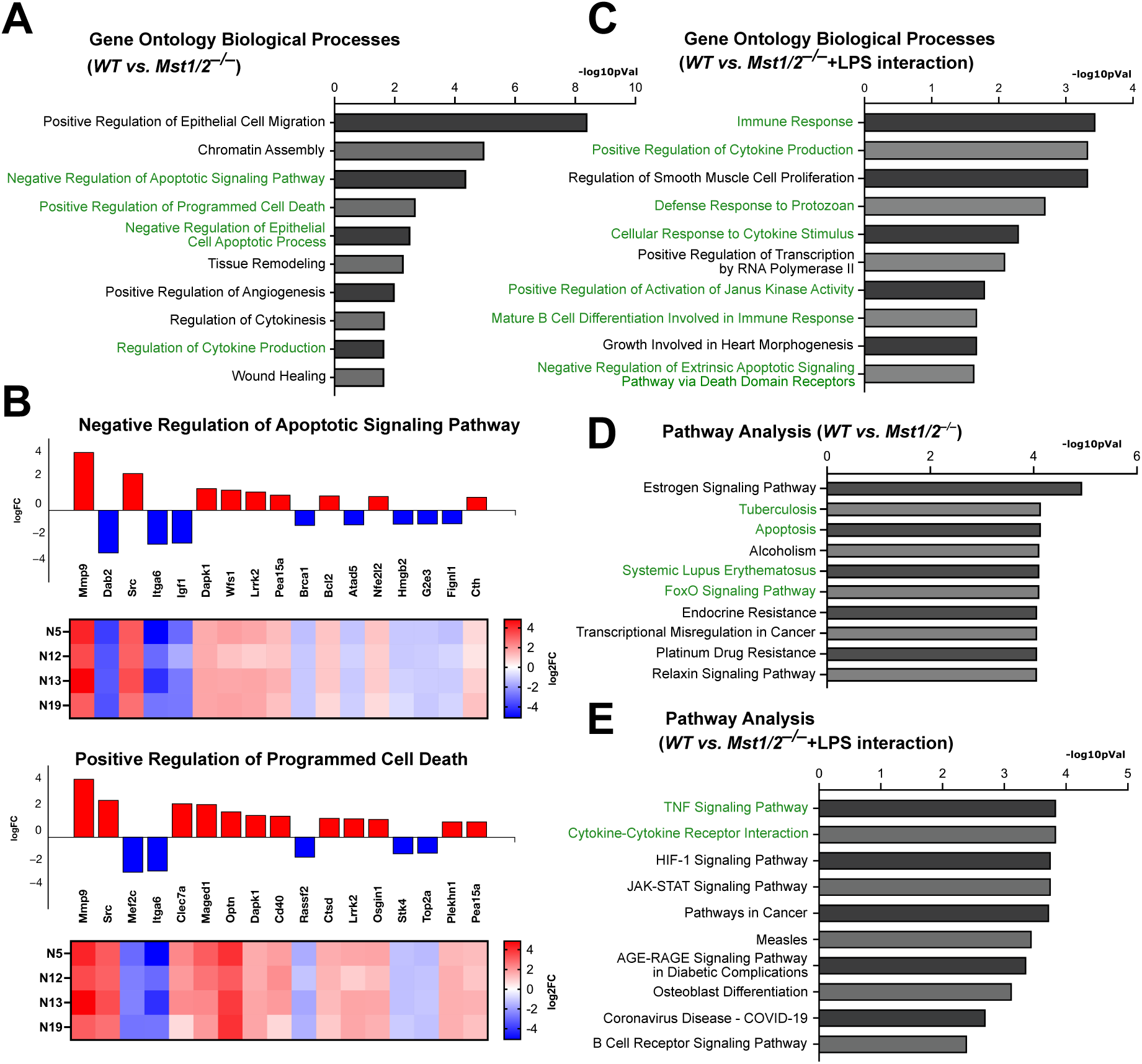
MST1/2 are the key regulators of immune pathways and biological processes in macrophages. **A.** Gene Ontology biological processes identified by the DEGs between WT and *Mst1/2^−/−^* iBMDMs. Up- and down-regulated significant gene profiles (360 significant genes) were combined between all *Mst1/2^−/−^* clones and provided to iPathwayGuide (AdvaitaBio). Processes are ranked by -log10 p value and smallest common denominator pruning is used for p-value correction. Top 10 processes with p values<0.05 were presented. Processes or pathways involved in immunity or cell death were in green fonts. **B.** Expression heat maps of four *Mst1/2^−/−^* clones for cell death relevant processes identified in A. Red, upregulated; blue, downregulated **C.** Gene Ontology biological processes identified from the 74 significant genes selected for the combined interaction effects of *Mst1/2* gene knockouts and LPS stimulation using iPathwayGuide. Processes are ranked by -log10 p value and smallest common denominator pruning is used for p-value correction. Top 10 processes with p values<0.05 were presented. **D.** KEGG Pathways identified by iPathwayGuide based on the 360 significant genes of *Mst1/2^−/−^* vs. WT iBMDMS. The false discovery rates (FDRs) were presented as -log10 p values and the top 10 pathways were presented. **E.** KEGG Pathways identified by iPathwayGuide based on the 74 significant genes selected for the combined interaction effects of *Mst1/2* gene knockouts and LPS stimulation. The FDRs were presented as -log10 p values and the top 10 pathways were presented.

### MST1/2 regulate cathepsin K (Ctsk) and B-cell lymphoma 2 (BCL2) expression via non-canonical Hippo signaling

RNAseq revealed that *Ctsk* encoding the lysosomal protease cathepsin K was abundantly transcribed in *Mst1/2^−/−^* macrophages (Fig. 1C and 3A) but barely detected in WT macrophages. CTSK is highly expressed in osteoclasts, a type of macrophages involved in bone resorption, and plays a role in autoimmune disorders, such as psoriasis and systemic lupus erythematosus (34, 35). Since regulation of CTSK by MST1/2 was not reported before, we confirmed that both the pro-form and mature CTSK proteins were highly produced in *Mst1/2^−/−^* macrophages but below detection in WT macrophages by immunoblotting (Fig. 3A). Given that BCL2 is a well-characterized regulator of apoptosis (36) and one of the DEG controlled by MST1/2 (Fig. 1C), we also determined the BCL2 protein expression. Consistent with the levels of RNA transcripts, increased BCL2 protein levels were detected in *Mst1/2^−/−^* macrophages (Fig. 3B). MST1 and MST2 are highly similar (Fig. S3) and phosphorylate the downstream Hippo components, such as MOB1 (6). To investigate whether MST1/2 have shared or distinct activities in gene expression, we generated iBMDMs expressing either MST1 (*Mst1+*) or MST2 (*Mst2+*) only. While CTSK was highly expressed in the double-knockout *Mst1/2^−/−^* macrophages, its protein level was suppressed in *Mst1+* or *Mst2+* macrophages (Fig. 3C). *Mst1+* macrophages, like WT macrophages, had low levels of BCL2, but *Mst2+* macrophages had increased BCL2 proteins comparable to the level in *Mst1/2^−/−^* macrophages (Fig. 3C), revealing that the two kinases have overlapping but also separate activities in gene expression.

**Figure 3.**
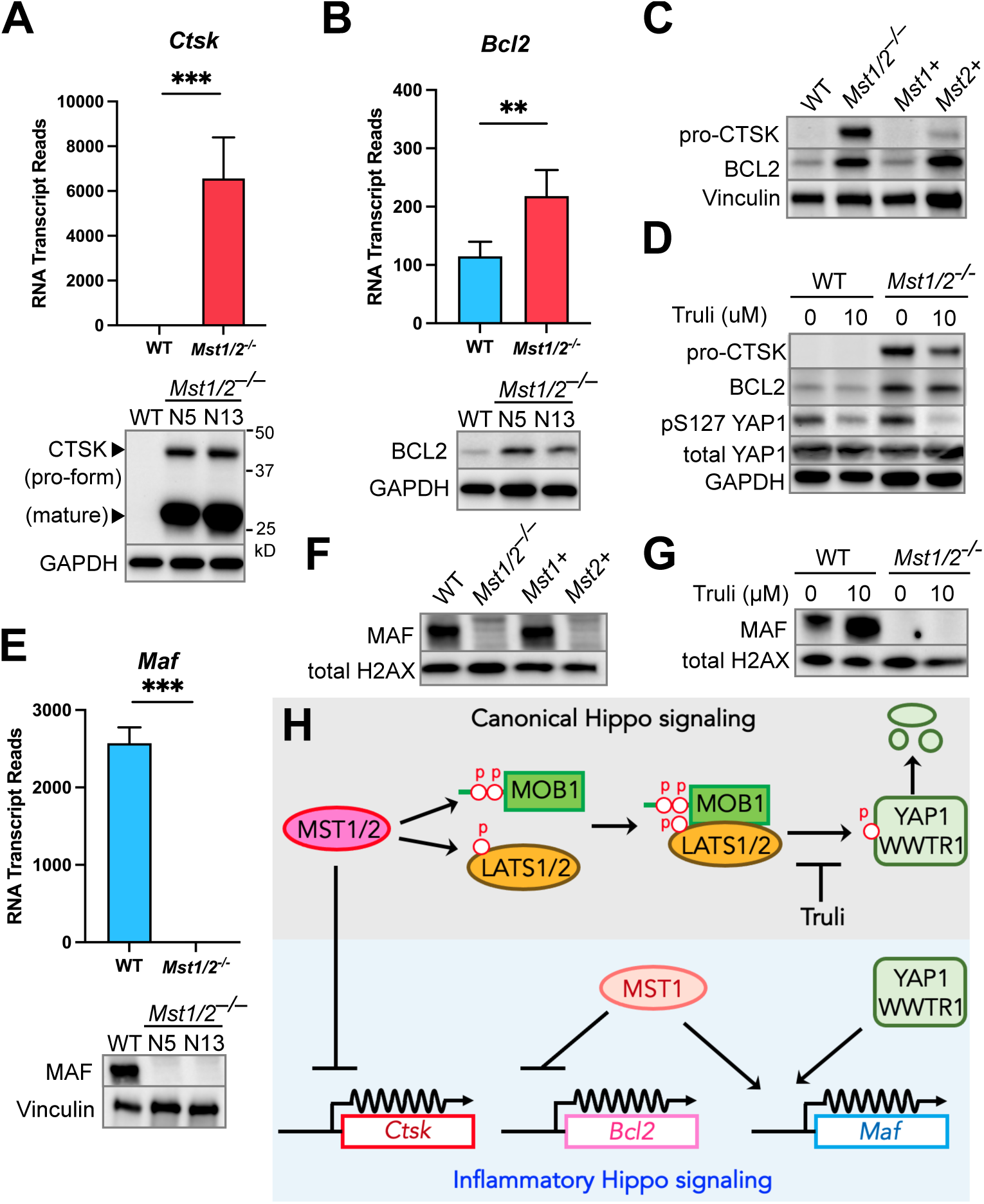
Inflammatory Hippo signaling for macrophage gene expression is independent of the canonical pathway. **A, B and E**. RNA and protein expression in WT and *Mst1/2^−/−^* iBMDMs. Top: Average RNA transcript reads for *Ctsk, Bcl2* and *Maf* from the RNAseq data of *Mst1/2^−/−^* N5 clone and WT iBMDMs. RNA average reads from five independent repeats. Student’s t-test, two tailed, unpaired, **: p < 0.01; ***: p <0.001. Bottom: Protein levels of CTSK, BCL2 and MAF in cell lysate of WT iBMDMs and two *Mst1/2^−/−^* iBMDM clones were detected by immunoblotting. **C and F**. Levels of the indicated proteins in WT, *Mst1/2^−/−^*, MST1-expressing (*Mst1+*), and MST2-expressing (*Mst2+*) iBMDMs were determined by immunoblots. **D and G**: WT or *Mst1/2^−/−^* iBMDMs were treated with or without 10 µM Truli for 24 hours. Cells were then lysed, and protein samples were collected and analyzed by immunoblots. Vinculin, GAPDH, and Total histone H2AX were used as internal loading controls in immunoblots. All blots are representative of three independent repeats. **H**. Schematic model of the regulatory activities of MST1 and MST2 through inflammatory Hippo signaling to control expression of *Ctsk*, *Bcl2* and *Maf* in macrophages.

When canonical Hippo signaling is on, MST1/2 phosphorylate and activate the downstream kinases LATS1/2 which phosphorylate YAP1/WWTR1 (Fig. 3H). Since CTSK and BCL2 expression was increased in *Mst1/2^−/−^* macrophages, we used a nucleotide-analog inhibitor, Truli (37), to specifically block the kinase activity of LATS1/2, resulting in an off status of canonical Hippo signaling that is similar to knocking out *Mst1/2*. As anticipated, Truli reduced YAP1 phosphorylation (Fig. 3D). No change in BCL2 protein expression was detected in WT or *Mst1/2^−/−^* macrophages treated with Truli (Fig. 3D). Truli also did not induce CTSK expression in WT macrophages and, unexpectedly, caused moderate decrease of CTSK in *Mst1/2^−/−^* macrophages (Fig. 3D), indicating expression of CTSK and BCL2 are controlled by MST1/2 via alternative routes other than canonical Hippo signaling.

### MST1 up-regulates Macrophage activating factor (MAF) expression through a mechanism that is not canonical Hippo signaling

We next characterized expression of *Maf*, one of the highly down-regulated genes in *Mst1/2^−/−^* macrophages (Fig. 1C). MAF is a leucine zipper transcription factor (38) and controls key functions in macrophages, including self-renewal, cell death and cytokine production (39–42). While both RNA and protein expression of *Maf* were detected in WT macrophages, lack of MST1/2 resulted in no detectable MAF (Fig. 3E). Unlike CTSK being negatively controlled by both MST1 and MST2, only *Mst1+* macrophages restored MAF expression (Fig. 3F). Interestingly, the Truli treatment led to an increase of MAF expression in WT macrophages while MAF remained below detection in *Mst1/2^−/−^* macrophages (Fig. 3G). This result reveals a unique regulatory loop that the presence of MST1 kinase positively regulates MAF expression, likely through a separate mechanism, and canonical Hippo signaling mediated by LATS1/2 kinases suppresses MAF expression. Together with CTSK and BCL2 regulation, these findings show that MST1/2 have shared and distinct activities in macrophage gene expression via signaling cascades independent of canonical Hippo signaling (Fig. 3H).

### MST1/2 affect release of cytokines and chemokines in macrophages

GO analyses identified several biological processes involved in cytokine production were potentially affected by MST1/2 (Fig. 2). Therefore, we sought to use cytokine arrays to detect the presence of 40 cytokines in the conditioned media collected from WT and *Mst1/2^−/−^* macrophages over 3 hours of incubation (Fig. S4A). The relative secretion level of CxC motif chemokine 10 (CXCL10) was significantly reduced in the conditioned media of *Mst1/2^−/−^* macrophages (Fig. 4A) while secretion of TNFα and interleukin-1 receptor antagonist protein (IL1ra) were enhanced but not statistically significant. We challenged the iBMDMs with the Gram-negative bacterial pathogen *Legionella pneumophila* to determine if MST1/2 affect host cytokine release during infection. Upon challenge of the virulent *L. pneumophila* strain *Lp02* for 3 hours, the relative levels of TNFα, IL-1ra and intercellular adhesion molecule 1 (ICAM1) were significantly higher in the conditioned media collected from *Mst1/2^−/−^* macrophages than from WT macrophages (Fig. 4B). The level of CXCL10 secretion remained significantly reduced in *Mst1/2^−/−^*macrophages challenged with *Lp02* (Fig. 4B). The results from the cytokine arrays show that MST1/2 have a role in secretion of a variety of cytokines and chemokines by macrophages.

**Figure 4.**
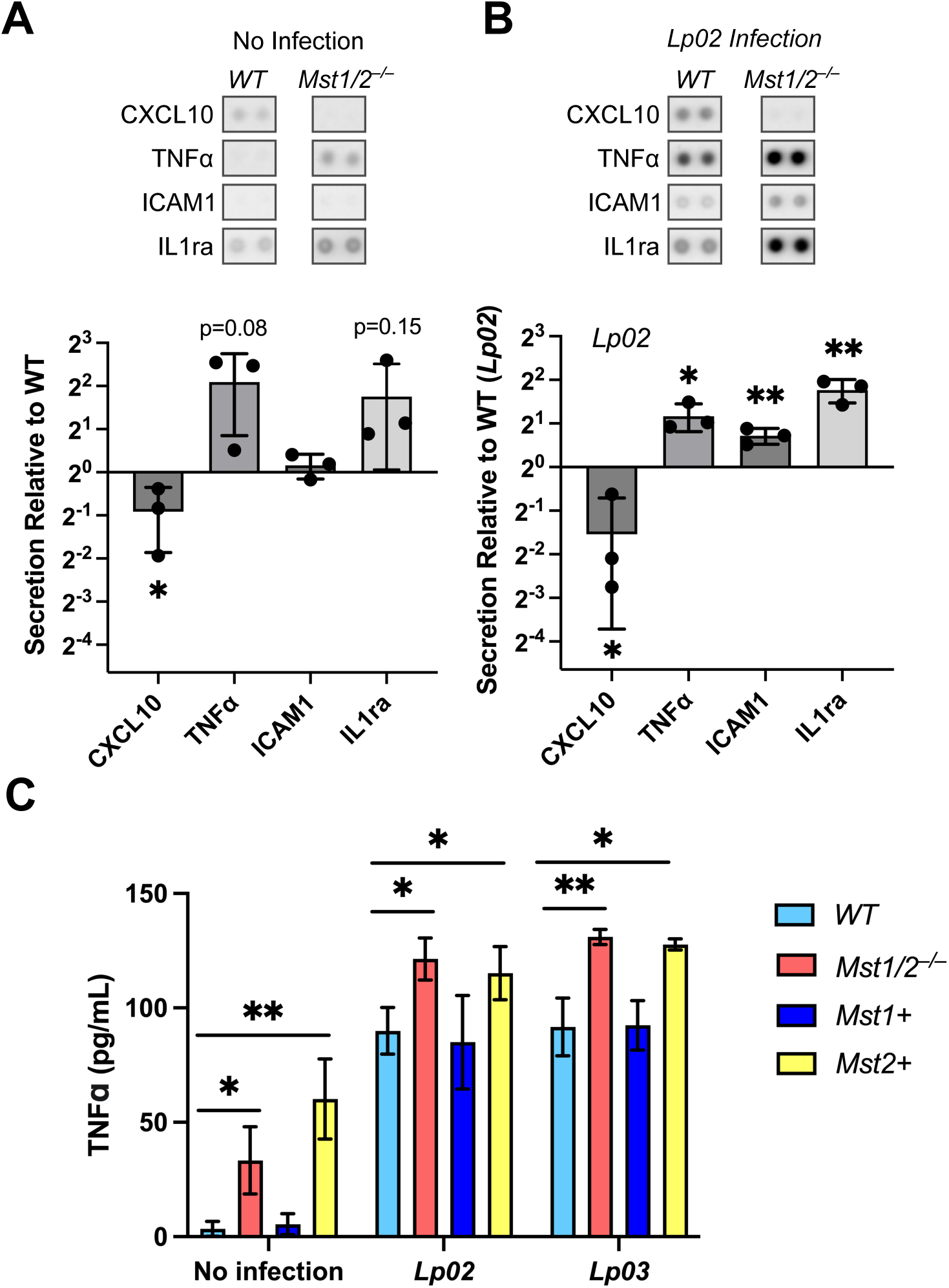
MST1/2 affect post-translational cytokine release in macrophages. **A and B**. Presence of cytokines and chemokines in the conditioned media collected from WT and *Mst1/2^−/−^* iBMDMs over 3 hours of incubation without infection (A) or with challenge of the *L. pneumophila Lp02* strain at MOI 10 (B) was detected by the cytokine arrays. Representative images of the cytokine spots on the arrays were presented. Graphs show relative secretion levels of the indicated cytokines in *Mst1/2^−/−^* and WT iBMDMs. The intensity of the cytokine spots was quantified using ImageJ and normalized to the intensity of the corresponding spots of WT iBMDMs (set as 1). Data were plotted as log2-fold change and presented as mean ± SD of three independent cytokine array experiments. **C**. Quantitation of TNFα secretion in WT, *Mst1/2^−/−^*, *Mst1+* and *Mst2+* BMDMs by ELISA. Conditioned media were collected from iBMDMs with or without challenge of the *L. pneumophila* strains (MOI=10) over 3 hours incubation. Concentrations of TNFα in the conditioned media were presented as mean ± SD (pg/mL) of three independent experiments. Student’s t-test, two tailed, unpaired, *: p < 0.05; **: p <0.01.

To determine whether MST1 and MST2 have different effects on cytokine secretion, we measured TNFα release by the *Mst1/2* double- and single-knockout iBMDMs using the quantitative enzyme-linked immunosorbent assays (ELISA). Confirming the cytokine array results, over 9-fold higher concentrations of TNFα were detected in the conditioned media of *Mst1/2^−/−^*macrophages than WT macrophages (average 33.3 pg/mL in *Mst1/2^−/−^*vs. 3.5 pg/mL in WT; Fig. 4C). *Mst1+* and WT macrophages released low levels of TNFα. Notably, *Mst2+* macrophages released more TNFα than WT and *Mst1+* macrophages at a level that was slightly and non-significantly higher than the level of *Mst1/2^−/−^* macrophages (Fig. 4C). Similar results were observed in conditioned media collected from the cells over 24 hours of incubation (Fig. S4B). We then compared TNFα release by the macrophages challenged with the virulent *L. pneumophila Lp02* strain or a non-virulent *Lp03* strain that lacks the type IV secretion system (T4SS). Challenging with either *Lp02* or *Lp03* stimulated release of TNFα by the macrophages. The TNFα levels in *Mst1/2^−/−^*and *Mst2+* macrophages remained significantly higher than in WT or *Mst1+* macrophages although the levels among these cells seemed to be relatively comparable (Fig. 4C). Thus, MST1 plays a suppressive role in TNFα secretion while the role of MST2 could be minor.

### MST1/2 promote apoptotic cell death in macrophages

The bioinformatic analyses pointed out that MST1/2 likely affect programmed cell death (Fig. 2), but the role of MST1/2 in macrophage cell death was undetermined. We tested activation of apoptosis in the iBMDMs with *Mst1/2* double- or single-knocked out. Treating WT macrophages with the apoptosis-inducing agent, staurosporine, triggered cleavage of poly-ADP-ribose polymerase-1 (PARP1) to an 89 kD fragment, a hallmark of apoptosis, and production of the activated apoptosis executioner caspase-3 (Casp3 p17) (Fig. 5A). In addition, we detected cleavage of full-length (FL) MST1/2 into the MST1/2 N-terminal (MST1/2-NT) fragments, a phenomenon that is also observed in several cell lines treated with staurosporine or Fas ligands (12–14, 43). In comparison to staurosporine that is a broad-spectrum kinase inhibitor, we treated the macrophages with Raptinal, a recently discovered agent that induces fast apoptosis via the intrinsic mitochondria pathway (44, 45). Raptinal induced production of MST1/2-NT and triggered apoptosis in WT macrophages (Fig. 5B). Importantly, when *Mst1/2^−/−^*macrophages were treated with low doses of staurosporine or Raptinal, PARP1 cleavage and Casp3 activation were reduced or minimal (Fig. 5). *Mst1+* or *Mst2+* macrophages triggered cleavage of MST1 or MST2, respectively, and had levels of apoptosis comparable to the level in WT iBMDMs, suggesting that MST1 or MST2 alone is sufficient to activate apoptosis induced by these treatments (Fig. 5). At high doses of staurosporine (2 μM) or Raptinal (10 μM), *Mst1/2^−/−^* macrophages underwent apoptosis (Fig. 5), indicating that excessive stimuli can bypass the requirement of MST1/2 to induce apoptosis. Overall, these finding demonstrate that MST1/2 are important modulators of the apoptotic death pathway in macrophages.

**Figure 5.**
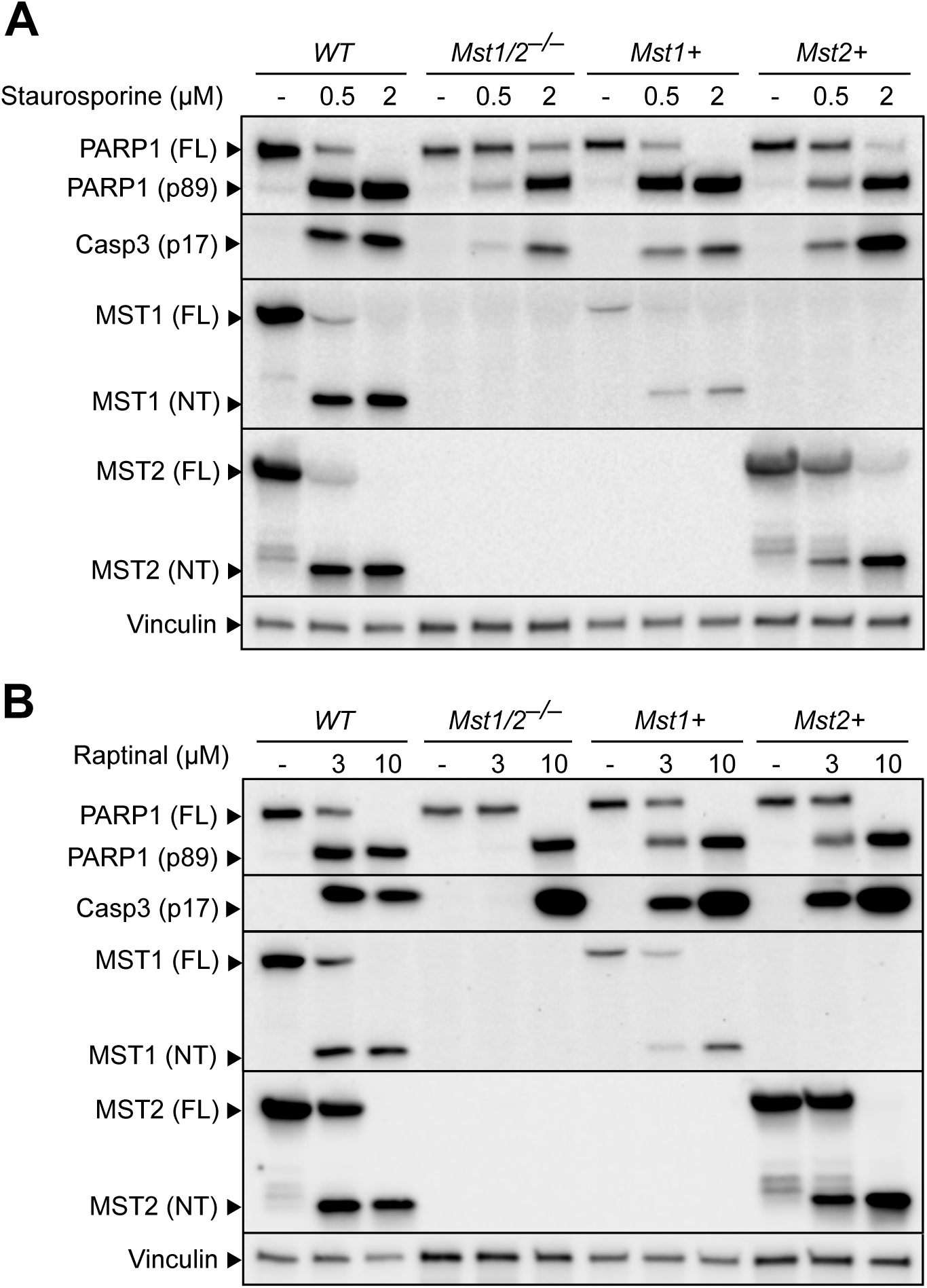
MST1/2 control activation of apoptosis in macrophages. Wildtype, *Mst1/2^−/−^, Mst1+, Mst2+* iBMDMs were treated with either staurosporine for 4 hours (A) or Raptinal for 3 hours (B) to induce apoptosis. Cell lysates were collected and levels of PARP1, MST1, MST2, activated caspase-3 (p17) and vinculin (as an internal control) were detected by immunoblotting.

### MST1/2 are the key factors in macrophages to restrict bacterial infection

Given that humans with MST1 deficiency experience recurrent infection (19, 20), we sought to determine whether lack of MST1/2 reduces macrophages’ ability to restrict invading bacteria. We challenged WT and *Mst1/2^−/−^* iBMDMs with a virulent *L. pneumophila* strain *Lp02ΔflaA* which had the flagellin gene deleted and measured its intracellular replication. The *Lp02ΔflaA* strain was used to avoid the interference from macrophage pyroptosis, a lytic form of programmed cell death, triggered by the presence of flagellin. Over the 72-hour infection period, *Lp02ΔflaA* had moderate intracellular growth in WT macrophages but grew robustly in *Mst1/2^−/−^* macrophages (Fig. 6A), suggesting that MST1/2 are required to restrict replication of *L. pneumophila*. To further elucidate individual role of MST1/2 in restricting bacterial infection, we challenged WT, *Mst1/2^−/−^*, *Mst1+* or *Mst2+* iBMDMs with *Escherichia coli* and performed a gentamicin protection assay. *Mst1/2^−/−^* macrophages had severe defects in restricting *E. coli* (Fig. 6B). Interestingly, *Mst2+* macrophages had increased susceptibility to *E. coli* although the level was significantly lower than the level in *Mst1/2^−/−^* macrophages (Fig. 6B). Expression of MST1 was sufficient to suppress *E. coli* infection. Similar results were obtained from the iBMDMs challenged with another bacterial pathogen *Pseudomonas aeruginosa* using the gentamicin protection assay (Fig. 6C). These results suggest that MST1 appears to play a major role in restriction of the bacteria and MST2 may have mild anti-bacterial activities.

**Figure 6.**
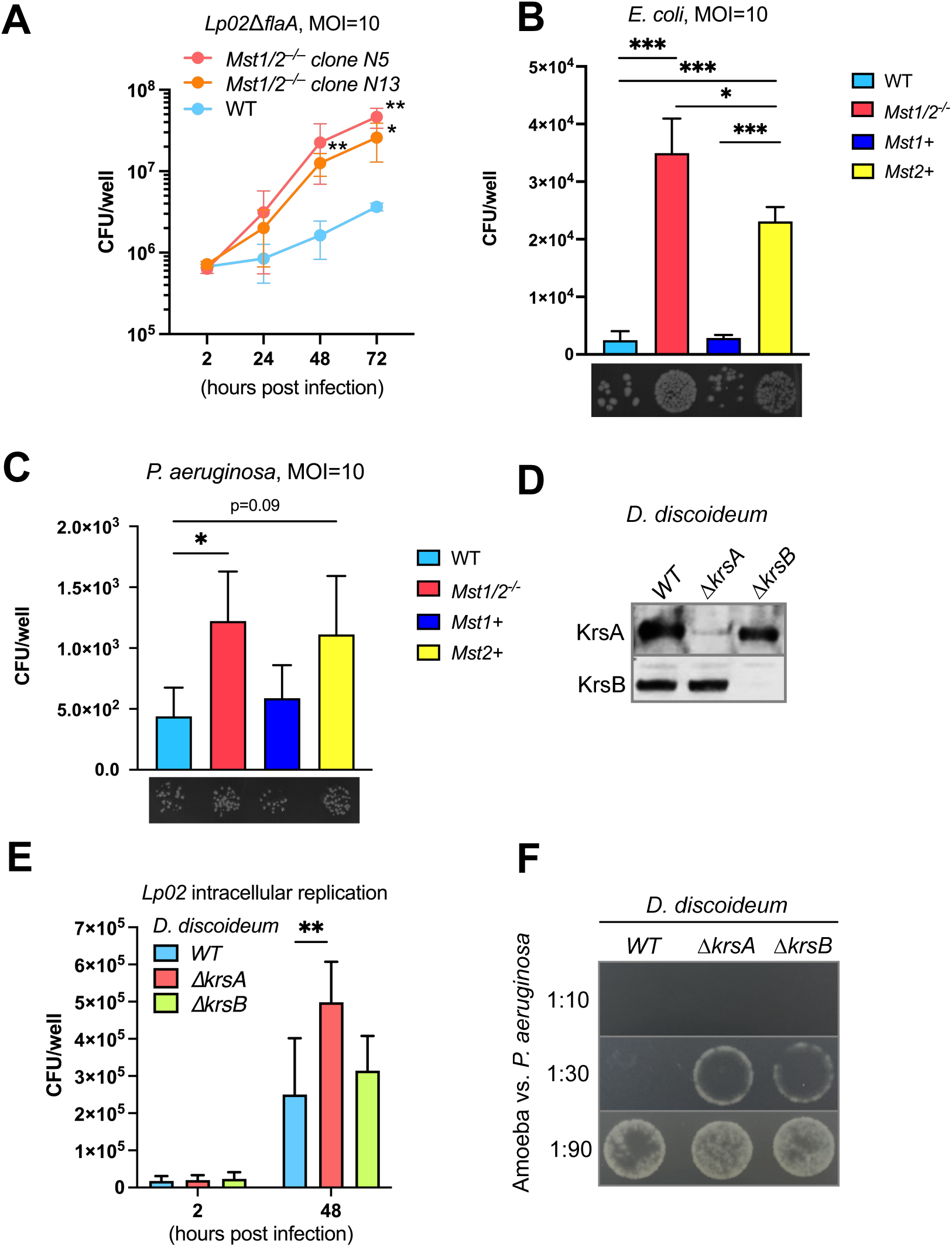
The conserved Hippo kinases in eukaryotic hosts restrict bacteria. **A.** WT and *Mst1/2^−/−^* iBMDMs were challenged with *L. pneumophila ΔflaA* at MOI=10. After 2 hours, free bacteria were removed by washing. At the indicated post infection time points, iBMDMs were lysed with digitonin to release intracellular bacteria. The numbers of *L. pneumophila* in the cell lysate were determined by the colony forming units (CFUs) assay. Data presented as mean± SD of three independent experiments. Student’s t-test, two tailed, unpaired, *: p < 0.05; **: p <0.01 compared to WT. **B and C.** iBMDMs were infected with *E. coli* or *P. aeruginosa* at MOI=10. After 1 hour, gentamicin (200 µg/mL) was added to kill extracellular bacteria. The infected iBMDMs were incubated for additional 2 hours and then lysed with Triton X-100. The cell lysate was serial diluted and plated on LB agar plates to count bacterial CFUs. Data presented as mean± SD of three independent experiments. Images represent formation of bacterial colonies on the plates. **D.** Protein levels of the Hippo kinases KrsA and KrsB in the *D. discoideum* strains were determined by immunoblotting. **E.** *D. discoideum* amoebae were challenged with the *L. pneumophila Lp02* strain at MOI=10. The numbers of *L. pneumophila* at the indicated post infection time points were determined by the CFUs assay as in A. Data presented as mean± SD of six independent experiments. **F**. Ability of the *D. discoideum* strains to prey on bacteria was determined by the semi-quantitation of resistance to predation. The amoebae were mixed with *P. aeruginosa* at the indicated ratios and the mixture was spot-platted on SM5 agar plates. Surviving bacteria grew on the plates after overnight incubation at 22℃. Images are representative of two independent experiments. Statistic method: Student’s t-test, two tailed, unpaired, *: p < 0.05; **: p <0.01.

### The Hippo kinases are conserved restriction factors against bacteria in protozoan hosts

*L. pneumophila* and *P. aeruginosa* are both opportunistic human pathogens and exist in natural environments. The Hippo pathway is highly conserved in eukaryotes, including free living protozoan that constantly encounter and prey on bacteria. We further investigated whether the Hippo kinase homologs, KrsA and KrsB (46) (Fig. S5), of the soil amoeba *Dictyostelium discoideum* affect the interactions with the bacteria. We obtained *D. discoideum* strains with *krsA* or *krsB* gene deletions and confirmed loss of KrsA or KrsB proteins in these strains (Fig. 6D). The *D. discoideum* strains were challenged with the *L. pneumophila* strain *Lp02* since *D. discoideum* cells do not undergo pyroptosis like macrophages. Because of the virulence factor T4SS, the *Lp02* strain was capable of replicating within *D. discoideum* and increased intracellular replication of *L. pneumophila* was observed in the *ΔkrsA* amoebae (Fig. 6E). We then used a semi-quantitative assay of resistance to predation to measure the ability of the *D. discoideum* strains in preying on and killing *P. aeruginosa*. In this assay, the amoebae and bacteria were mixed at different ratios and spotted on the SM5 agar plates to allow bacteria predation by amoebae. If amoebae failed to ingest and kill bacteria, surviving bacteria grew on the plates (47). As shown in Fig. 6F, the *ΔkrsA* and *ΔkrsB* strains were defective in predation or killing of *P. aeruginosa*. Therefore, the Hippo kinases also play a critical role in host defense in amoebae.

## Discussion

The Hippo kinases MST1/2 phosphorylate MOB1 and LATS1/2 to modulate transcriptional activity of YAP1/WWTR1, which is well-known as the canonical Hippo pathway (2, 3). Here, we investigate the diverse activities of the Hippo kinases in gene expression, release of cytokines/chemokines, programmed cell death, and antimicrobial defense in macrophages. Importantly, while being homologous proteins, MST1 and MST2 share certain functions but also individually possess unique activities. For example, both MST1/2 control macrophage apoptosis but have different effects on TNFα cytokine release (Fig. 5 and 4C). Moreover, we demonstrate that the Hippo kinases control anti-microbial activities in macrophages and amoebae, reflecting a conserved role of these kinases in determining host susceptibility to infection.

Since the datasets available for the transcriptome controlled by MST1/2 are mainly collected in non-immune cells, we profiled global gene expression of the macrophages lacking both MST1 and MST2. The MST1/2 transcriptome reveals genes that are previously not known to be regulated by MST1/2, such as *CtsK*, *Bcl2* and *Maf*, implying that MST1/2 control macrophage gene expression via an alternative signaling cascade (Fig. 1C and Fig. 3). This is also supported by the fact that most of the YAP1/WWTR1 target genes in non-immune cells are poorly expressed in macrophages (Fig. S1). Indeed, if CTSK and BCL2 expression were increased due to shutting off canonical Hippo signaling in *Mst1/2^−/−^* macrophages, blocking the activity of LATS1/2 by Truli would result in increased expression of CTSK or BCL2 in WT macrophages. In fact, the Truli treatment has no effect on CTSK or BCL2 expression (Fig. 3D). Likewise, Truli treatment promotes MAF expression in WT macrophages, an effect opposite to the suppressed expression observed in *Mst1/2^−/−^* macrophages (Fig. 3G). Taken together, MST1/2 regulate these representative genes in macrophages by alternative signaling cascades which we term the inflammatory Hippo pathway that are distinct from the canonical Hippo pathway (Fig. 3H).

The transcriptome analyses indicate importance of MST1/2 in immune pathways (Fig. 2). The cytokine arrays and ELISAs confirm the effects of MST1/2 on release of several cytokines and chemokines (Fig. 4). Although the effects of alteration in cytokine/chemokine release on immune cell communication remain to be determined, these results indicate that post-translational protein trafficking is under control of the Hippo kinases in macrophages. In addition, MST1 and MST2 have different activities in regulating TNFα release by macrophages (Fig. 4C). Challenging with *L. pneumophila* markedly increased TNFα release in all of the macrophage clones, and similar levels of TNFα were detected between macrophages challenged with the virulent T4SS-expressing *Lp02* strain and the non-virulent T4SS-null *Lp03* strain, suggesting that *L. pneumophila*-induced TNFα release is independent of the virulence factor T4SS. Similar effects on TNFα release have been observed in macrophages challenged with live or heat-killed *L. pneumophila* (48). Thus, MST1, but not MST2, has a role in preventing premature TNFα release in macrophages prior to encountering invading bacteria.

Both MST1 and MST2 can phosphorylate MOB1 and LATS1/2 to activate the canonical Hippo pathway (Fig. 3H). Expression of either MST1 or MST2 markedly reduces CTSK expression (Fig. 3C) although the results from the Truli experiment suggests that the suppression is mediated by an alternative, non-canonical Hippo signaling. On the other hand, like their different roles in TNFα release, we observed that MST1 has significant impacts on BCL2 and MAF expression while MST2 has no effect (Fig. 3C and 3F). The differences between MST1 and MST2 in gene regulation further demonstrate that these Hippo kinases affect immunity independent of the canonical Hippo pathway. Indeed, MST1/2 can affect autophagy by phosphorylating LC3, and the LATS1/2 kinases can be activated by an alternative route through MAP4Ks without MST1/2 (49–51).

Our results reveal the implications of the inflammatory Hippo pathway in macrophage differentiation and autoimmune diseases. For example, MST1 is essential for expression of MAF which has a role in cell death, inflammatory responses of the tumor-associated macrophages, and macrophage differentiation (40–42, 52). MST1/2 strongly suppress expression of CTSK that is highly produced in osteoclasts, the bone macrophages, and has been a therapeutic target for treating osteoporosis (53). CTSK also contributes to pathogenesis of systemic lupus erythematosus (SLE) (34), one of the top pathways identified in the bioinformatic analyses (Fig. 2C). This is correlated with expression of the complement genes *C1qa, b, c* being highly suppressed in *Mst1/2^−/−^*macrophages (Fig. 1C, 1D). Although the *C1q* genes were not selected for the iPathwayGuide analysis due to the stringent criteria used, C1q deficiency is a pathogenic factor of SLE (54, 55). Collectively, identifying MST1/2 as key regulators of these genes offers insights to manage anti-tumor immunity or mitigate osteoporosis and SLE.

Programmed cell death is an important host defense in innate immunity (56). The pro-apoptotic activity of MST1/2 has been characterized in cancer cell lines or fibroblasts by overproducing the kinases or gene knockdown methods (12–16). In contrast, *Mst1/2^−/−^* naïve T cells are prone to apoptosis (18–20). We found that MST1 and MST2 are both required for macrophage apoptosis with low-dose treatments of staurosporine and Raptinal but can be bypassed if the cells are over-stimulated at high doses (Fig. 5), indicating that macrophages with *Mst1/2* knockouts still possess the essential components of the apoptotic death pathway and the gene knockouts did not cause secondary mutations in the apoptotic cascade. BCL2 has anti-apoptosis activity by preventing activation of the intrinsic pathway. While *Mst1/2^−/−^* and *Mst2+* macrophages express increased BCL2 (Fig. 3C), *Mst2+* macrophages still undergo apoptosis when treated with the apoptosis agents at low doses (Fig. 5), suggesting that the resistance to apoptosis in *Mst1/2^−/−^* macrophages is likely caused by other mechanisms. The C-termini of MST1/2 have self-inhibitory activity, and removal of the C-termini by proteases increases the kinase activity of MST1/2, resulting in activation of the apoptosis (57). Therefore, MST1/2-NT produced upon the treatments may lead to apoptosis in macrophages (Fig.5). Notably, studies showed that the apoptotic caspase Casp3 cleaves MST1/2 and produces MST1/2-NT to promote apoptosis in non-macrophage cell lines (12, 14). In this study, *Mst1/2* knockout macrophages have reduced Casp3 activation in response to the low-dose treatments (Fig. 5). These observations suggest that MST1/2 can regulate Casp3 activation and functions as modulators of apoptosis in macrophages.

The results from the infection experiments (Fig. 6) demonstrate the critical role of MST1/2 in anti-bacterial host defense, showing a correlation with the increased susceptibility to infection observed in humans with MST1 deficiency and *Mst1/2* double knockout mice (19, 20, 23). *Mst1/2^−/−^* macrophages fail to limit infections by *E. coli*, *P. aeruginosa* and *L. pneumophila* (Fig. 6A to C). We found that MST1 has a more pivotal role in anti-bacterial activities than MST2 (Fig. 6B, 6C) and that MST1/2 may activate different host defense mechanisms against different bacteria. For example, expression of MST2 in macrophages partially restores the anti-bacterial activity against *E. coli* but not *P. aeruginosa* (Fig. 6B, 6C). Similarly, *ΔkrsA* amoebae are more permissive to the intracellular bacterium *L. pneumophila* while either *ΔkrsA* or *ΔkrsB* amoebae are incompetent in preying and killing the extracellular bacterium *P. aeruginosa* (Fig. 6E, 6F). This indicates that the single Hippo kinase knockout *D. discoideum* also have different types of defects in host defense. Collectively, the Hippo kinases are key factors in controlling a range of host defense mechanisms in mammalian macrophages and protozoans upon the types of pathogen challenge. Future studies that dissect these mechanisms in the host cells at molecular levels are necessary to understand the central role of these conserved Hippo kinases in host-pathogen interactions and innate immunity.

## Data availability

RNA sequencing raw data are available in fastq format and deposited in (to be determined). Other data are presented in this manuscript or available in the supplementary materials.

## Acknowledgements

This study is supported by Wayne State University Startup Funds (to P.L.), the University Research Grant 2020-21, Wayne State University (to P.L.), and NIH grant number P30 CA022453 (to G.D.). We thank Dr. Yulia Artemenko (State University of New York, Oswego) for sharing the *D. discoideum* strains and antibodies against KrsA and KrsB. Author contribution: Study design-P.L., B.M.S., S.M.Q.; Perform experiments and collect data-P.L., B.M.S., S.M.Q, Y.S.; Analyze data-P.L., B.M.S., S.M.Q., G.D.; Write manuscript-P.L., B.M.S., S.M.Q.

## Materials and Methods

### Immortalized bone marrow-derived macrophages culture

Mouse immortalized bone marrow derived macrophages (iBMDMs) were gifts from Dr. Jonathan Kagan. Cells were cultured in Dulbecco’s Modified Eagle’s medium (DMEM, Corning 15013CV) supplemented with 10% fetal bovine serum (FBS, Corning 35-016-CV), 2mM L-glutamine (Corning, MT25005CI) and 0.1% penicillin/streptomycin (P/S, Corning, MT3000CI) at 37°C in humidified incubators with 5% CO. Cells were passaged every 2-3 days when they reached 60-80% confluency.

### Knocking out Mst1 and Mst2 genes in iBMDMs

LentiCas9-Blast (Addgene #52962) plasmid containing human codon-optimized Streptococcus pyogenes Cas9 protein and Stk3 (Mst2, Addgene #75975) gRNA was used to dually target *Mst1* and *Mst2*, as the gRNA sequence is identical in both genes, and packaged in lentiviral particles. Lentiviral particles were generated by co-transfection with pMDLg/RRE (Addgene #12251) and pMD2.G (Addgene #12259, gifted to us by Didier Trono) into HEK293T/17 cells for 24 hours. Following transfection, culture supernatant was collected, centrifuged, and filtered. Culture supernatants were added to parental iBMDMs to undergo a 72-hour transduction. Following the transduction, iBMDMs were selected using 10 µg/mL blasticidin. For *Mst1* and *Mst2* double or single knockouts, Cas9 expressing iBMDMs were transduced for 72 hours with viral particles containing Stk3 gRNA. Following transduction, iBMDMs were selected by using 10 µg/mL blasticidin and 10 µg/mL puromycin. Blasticidin and puromycin-resistant iBMDMs were isolated through limited dilutions and amplification. Cells were then lysed using 1X Laemmli sample buffer and proteins were analyzed by immunoblotting to examine MST1 and MST2 expression. Cas9-expressing iBMDMs were used as the wildtype control in all following experiments.

### Bacterial culture

*E. coli* strain MG1655 was suspended in 2.5mL of regular LB broth (10 g/L tryptone, 5 g/L yeast extract, 10 g/L NaCl,) and incubated overnight at 37°C in a shaking incubator. *Pseudomonas aeruginosa* strain PAO1F was suspended 2.5mL in liquid high salt LB broth (10 g/L tryptone, 5 g/L yeast extract, 11.7 g/L NaCl, 10 mM MgCl_2_, 0.5 mM CaCl_2_) and incubated overnight at 37°C in a shaking incubator. The next day, the bacterial cultures were diluted at 1:50 or 1:100 in fresh LB broth (for E. coli) or high salt LB (for *P. aeruginosa*) and incubated for 2-3 hours at 37°C in a shaking incubator. The fresh bacterial culture was pelleted at 11,000xg for 2 minutes. Bacterial pellets were resuspended in DMEM without FBS or antibiotics. Density of bacterial resuspension was measured at optical density (OD600) and used to calculate the dilutions for the desired multiplicity of infection (MOI).

*Legionella pneumophila* patch cultures on CAYE agar plate (2 g/L activated charcoal, 10 g/L ACES, 10 g/L yeast extract, 400 µg/mL cysteine, 135 µg/mL ferric nitrate, 50 µg/mL streptomycin, 15g/L agar) were resuspended in liquid AYE broth (10g/L ACES, 10g/L yeast extract, 400µg/mL cysteine, 135µg/mL ferric nitrate) and incubated at 37°C. Overnight cultures were centrifuged at 11,000xg for 2 minutes, then resuspended in DMEM supplemented with 10% FBS and 2mM L-glutamine without antibiotics. Density of bacterial resuspension was measured at optical density (OD600) and used to calculate the dilutions for the desired multiplicity of infection (MOI).

### Dictyostelium discoideum culture

*D. discoideum*, including the wildtype AX2 parental strain (strain ID: DBS0350762), ΔkrsA strain (strain ID: DBS0350759) and ΔkrsB strain (strain ID: DBS0350760), were obtained from the DictyBase stock center (Northwestern University, Chicago, USA). Axenic *D. discoideum* cells were cultured in HL-5 media (14 g/L glucose, 7 g/L yeast extract, 14 g/L thiotone, 0.95 g/L Na_2_HPO_4_-7H_2_O, 0.5 g/L KH_2_PO_4_, pH 6.5) at 21°C in a microbiological incubator. The amoeba cells were sub-cultured every two days. For measuring *L. pneumophila* intracellular replication, the amoebae were harvest and resuspended in MB media (3.9 g/L MES, 7 g/L yeast extract, 14 g/L thiotone, pH6.9). 5x10^5^ *D. discoideum*/well were seeded in 24-well plates and incubate at 26 °C for two hours. 50 µL of *L. pneumophila* suspension in MB media was added into the wells to challenge the *D. discoideum* at MOI=10. The plates were centrifuged at 200xg for 5 minutes and incubated at 26°C for additional two hours. After the incubation, the wells were washed with fresh MB media three times to remove extracellular *L. pneumophila*. 0.5 mL/well of fresh MB media was added into the wells and the plates were incubated at 26°C in a microbiological incubator. At the desired post infection timepoints, infected *D. discoideum* cells were lysed with sodium deoxycholate (final concentration 0.05%) to release intracellular *L. pneumophila*. The lysate was serial diluted with sterile water and spot platted on CAYE agar plates, and the numbers of *L. pneumophila* were determined by colony forming units (CFUs) assays.

### Gentamicin protection assays

iBMDMs (1x10^5^/well) were seeded into 24-well plates. The next day, media was removed and replaced with fresh DMEM without FBS or P/S. iBMDMs were challenged with *E. coli* or *P. aeruginosa* and centrifuged at 200 xg for 5 minutes to facilitate contact between bacteria and iBMDMs. Cells were incubated at 37°C for 1 hour followed by adding 200 µg/mL gentamicin. Cells were incubated for an additional 2 hours, then washed 3 times with DMEM and lysed with 500 µL of 0.1% Triton-X100 in sterile water per well for 10 minutes. The lysate was serial diluted and spotted onto LB agar plates. Plates were incubated overnight at 30°C and colonies were counted the following day to determine colony forming units (CFUs).

### Semi-quantitation assays of resistance to predation of D. discoideum

*D. discoideum* cells cultured in MB media were harvested and resuspended in SorC buffer (2 g/L KH_2_PO_4_, 0.55 g Na_2_HPO_4_-7H_2_O, 50 µM CaCl_2_) at 5x10^6^, 1.675x10^6^, 0.56x10^6^ cells/mL (3-fold serial dilutions). Overnight *P. aeruginosa* cultures were 1:100 diluted in high salt LB and cultured at 37 °C for 2-3 hours to reach mid-log growth phase. *P. aeruginosa* was pelleted and resuspend in SorC buffer at 5x10^7^ bacteria/mL. 100 µL (5x10^6^) of *P. aeruginosa* were mixed with 100 µL of the serial diluted *D. discoideum* to have the bacteria: amoeba ratios at 10:1, 30:1, and 90:1. The amoeba/bacteria mixtures were spot platted on SM5 agar plates (2 g/L glucose, 2 g/L bactopeptone, 2 g/mL yeast extract, 0.2 g/L MgSO_4_-7H_2_O, 1.9 g/L KH_2_PO_4_, 1g/L K_2_HPO_4_, 20 g/L agar). The SM5 agar plates were incubated at 22°C in a microbiological incubator and imaged the next day.

### L. pneumophila intracellular replication in iBMDMs

WT and *Mst1/2^−/−^* iBMDMs were seeded at 1x10^5^ cells/well in 24-well plates one day prior to infection. *L. pneumophila* with an in-frame deletion of the flagellin gene *flaA* (Lp02ΔflaA) overnight cultures were pelleted and resuspended in fresh DMEM without P/S. The culture media in the 24-well plates were replaced with 900 µL of fresh DMEM without P/S, and 100 µL of *L. pneumophila* suspension was added to the wells to challenge the macrophages at MOI=10. The plates were centrifuged at 200xg for 5 minutes and incubated in the cell culture incubator. After two hours of incubation, the wells were washed with 1xPBS for three times to remove free bacteria, and 1 mL of fresh DMEM without P/S was added. The plates were incubated in a cell culture incubator to the indicated post infection timepoints. To count the numbers of *L. pneumophila* at the indicated time points, iBMDMs were lysed with 0.01% digitonin to release intracellular *L. pneumophila*. The lysate was serial diluted with sterile water and spot plated on CAYE agar plates. The agar plates were incubated at 37°C in a microbiological incubator for 4 days and the colony forming units were calculated.

### RNA Sequencing

WT and *Mst1/2^−/−^* iBMDMs were seeded in 6-well plates at 3x10^5^ cells/well. After culturing for 48 hours, the media was removed and replaced with fresh DMEM with or without LPS (2 µg/mL) for 3 hours. Total RNA samples from five independent repeats were extracted using a Qiagen RNAeasy mini kit (Cat. #74104) according to the manufacturer’s manual. Extracted and isolated RNA was sent to the Genomics Core at Wayne State University for sequencing and analysis using a QuantSeq 3’ mRNA-Seq Library Prep Kit. RNA samples were sent to the Genomics Core at Wayne State University for sequencing and analyses. RNA quality control and quantification was performed on the TapeStation 4100, and libraries were prepared with 100 ng input using the QuantSeq 3’ mRNA-seq Library Prep Kit. Prepared RNA libraries were sequenced on the NovaSeq 6000 (>10M reads per sample). Reads were aligned to the mouse genome (mm10) with STAR for alignment followed by ht-seq count for tabulation for each gene region. Differential expression was analyzed with the log2 fold change between conditions, p-value, false discovery rate corrected p-value, and individual read counts.

### Pathway and Gene Ontology biological process analyses

RNA sequencing read counts were transformed by multiplying by 1,000,000 and dividing by the total number of reads for each sample to obtain counts per million reads mapped (CPM), which was then log2 transformed (after adding a small positive constant) to account for the inherent skewness of RNAseq count data. For each measured gene, the optimal statistical model describing the relationship between *Mst1/2* double knockout status and LPS treatment with CPM was chosen using the Bayesian Information Criterion. These investigated models included main effect only and interaction models. Bioinformatics analyses were then conducted using iPathwayGuide (Advaita) including and excluding interaction effects with a reduced gene pool of those with a ± log2-fold change, p-value < 0.05, and an overall mean of at least 2 on the CPM scale.

### Cytokine arrays and enzyme-linked immunosorbent assays (ELISA)

iBMDMs were seeded in 12-well plates (3x10^5^/well). The following day, media was removed and replaced with 1 mL of fresh DMEM, and the cells were incubated at 37°C for 3 or 24 hours. The culture media were centrifuged at 200xg for 5 minutes and 800 µL of supernatant was collected. The supernatant was centrifuged again at 2,000xg for 10 minutes to remove cell debris. The supernatant after the second centrifugation was analyzed by the cytokine arrays kits (R&D systems, ARY006) or the TNFα ELISA kit (BioLegend, #430904) according to the procedures described in the manufacturers’ manuals.

### Truli, Raptinal, and staurosporine treatment

iBMDMs (3x10^5^ cells/well) were seeded one day prior to experimentation in 12-well plates. Prior to the treatments, media was removed and replaced with fresh DMEM. iBMDMs were treated with DMSO vehicle control, Truli (N-(3-Benzylthiazol-2(3H)-ylidene)-1H-pyrrolo[2,3-b] pyridine-3-carboxamide; CSN pharm CSN26140) or Raptinal (Adipogene, AG-CR1-2902) and incubated in CO2 humidified 37°C incubators for 3 hours. For staurosporine experiments, iBMDMs were treated with DMSO vehicle control or staurosporine (Cayman chemical, 81590) for 4 hours. Following the treatments, cell culture media was transferred to microcentrifuge tubes and centrifuged 200xg for 5 minutes to obtain a cell pellet. During centrifugation, cell monolayers in 12-well plates were lysed with 1X Laemmli buffer. Following centrifugation, supernatant was removed, and remaining cell pellet was combined with respective cell lysate from the treatments. Cell lysate was denatured at 97°C for 10 minutes and analyzed by SDS-PAGE and immunoblotting. Immunoblots were developed by horse-radish peroxidase (HRP) chemiluminescence and imaged using the BioRad Chemidoc Imaging system. Antibodies used: vinculin (sc-73614), GAPDH (sc-25778), BCL2 (sc-7382), and CTSK (sc-48353) from Santa Cruz Biotechnology. MST2 (ab52641) from Abcam. MST1 (14946S), cleaved caspase-3 (9661S), PARP1 (9542S), phospho-S139 H2AX (2577S), total H2AX (2595S), MOB1 (13730S), phospho-T35 MOB1 (8699S), and LATS1 (3477S) from Cell Signaling. MAF (A300-613A) from Bethyl Laboratories. Rabbit polyclonal antibodies against *D. discoideum* KrsA and KrsB were gifts from Dr. Yulia Artemenko (State University of New York, Oswego) (46). All immunoblots are representative of at least three biological repeats.

### Statistical Analyses and Graphing

Statistical analyses and graphs were performed and produced in GraphPad Prism and R.

## References

1. Hilman D, Gat U. 2011. The Evolutionary History of YAP and the Hippo/YAP Pathway. Molecular Biology and Evolution 28:2403–2417.

2. Yu F-X, Zhao B, Guan K-L. 2015. Hippo Pathway in Organ Size Control, Tissue Homeostasis, and Cancer. Cell 163:811–828.

3. Zheng Y, Pan D. 2019. The Hippo Signaling Pathway in Development and Disease. Dev Cell 50:264–282.

4. Dong J, Feldmann G, Huang J, Wu S, Zhang N, Comerford SA, Gayyed MF, Anders RA, Maitra A, Pan D. 2007. Elucidation of a Universal Size-Control Mechanism in Drosophila and Mammals. Cell 130:1120–1133.

5. Zhao B, Ye X, Yu J, Li L, Li W, Li S, Yu J, Lin JD, Wang C-Y, Chinnaiyan AM, Lai Z-C, Guan K-L. 2008. TEAD mediates YAP-dependent gene induction and growth control. Genes Dev 22:1962–1971.

6. Praskova M, Xia F, Avruch J. 2008. MOBKL1A/MOBKL1B Phosphorylation by MST1 and MST2 Inhibits Cell Proliferation. Current Biology 18:311–321.

7. Oh S, Lee D, Kim T, Kim T-S, Oh HJ, Hwang CY, Kong Y-Y, Kwon K-S, Lim D-S. 2009. Crucial Role for Mst1 and Mst2 Kinases in Early Embryonic Development of the Mouse. Mol Cell Biol 29:6309–6320.

8. Zhou D, Conrad C, Xia F, Park J-S, Payer B, Yin Y, Lauwers GY, Thasler W, Lee JT, Avruch J, Bardeesy N. 2009. Mst1 and Mst2 maintain hepatocyte quiescence and suppress hepatocellular carcinoma development through inactivation of the Yap1 oncogene. Cancer Cell 16:425–438.

9. Song H, Mak KK, Topol L, Yun K, Hu J, Garrett L, Chen Y, Park O, Chang J, Simpson RM, Wang C-Y, Gao B, Jiang J, Yang Y. 2010. Mammalian Mst1 and Mst2 kinases play essential roles in organ size control and tumor suppression. Proceedings of the National Academy of Sciences 107:1431–1436.

10. Lu L, Li Y, Kim SM, Bossuyt W, Liu P, Qiu Q, Wang Y, Halder G, Finegold MJ, Lee J-S, Johnson RL. 2010. Hippo signaling is a potent in vivo growth and tumor suppressor pathway in the mammalian liver. Proc Natl Acad Sci U S A 107:1437–1442.

11. Zhou D, Zhang Y, Wu H, Barry E, Yin Y, Lawrence E, Dawson D, Willis JE, Markowitz SD, Camargo FD, Avruch J. 2011. Mst1 and Mst2 protein kinases restrain intestinal stem cell proliferation and colonic tumorigenesis by inhibition of Yes-associated protein (Yap) overabundance. Proc Natl Acad Sci U S A 108:E1312–1320.

12. Graves JD, Gotoh Y, Draves KE, Ambrose D, Han DK, Wright M, Chernoff J, Clark EA, Krebs EG. 1998. Caspase-mediated activation and induction of apoptosis by the mammalian Ste20-like kinase Mst1. EMBO J 17:2224–2234.

13. Lee K-K, Ohyama T, Yajima N, Tsubuki S, Yonehara S. 2001. MST, a Physiological Caspase Substrate, Highly Sensitizes Apoptosis Both Upstream and Downstream of Caspase Activation*. Journal of Biological Chemistry 276:19276–19285.

14. Ura S, Masuyama N, Graves JD, Gotoh Y. 2001. Caspase cleavage of MST1 promotes nuclear translocation and chromatin condensation. Proceedings of the National Academy of Sciences 98:10148–10153.

15. Harvey KF, Pfleger CM, Hariharan IK. 2003. The Drosophila Mst Ortholog, hippo, Restricts Growth and Cell Proliferation and Promotes Apoptosis. Cell 114:457–467.

16. O’Neill E, Rushworth L, Baccarini M, Kolch W. 2004. Role of the Kinase MST2 in Suppression of Apoptosis by the Proto-Oncogene Product Raf-1. Science 306:2267–2270.

17. Steinhardt AA, Gayyed MF, Klein AP, Dong J, Maitra A, Pan D, Montgomery EA, Anders RA. 2008. Expression of Yes-associated protein in common solid tumors. Human Pathology 39:1582–1589.

18. Zhou D, Medoff BD, Chen L, Li L, Zhang X, Praskova M, Liu M, Landry A, Blumberg RS, Boussiotis VA, Xavier R, Avruch J. 2008. The Nore1B/Mst1 complex restrains antigen receptor-induced proliferation of naïve T cells. Proceedings of the National Academy of Sciences 105:20321–20326.

19. Abdollahpour H, Appaswamy G, Kotlarz D, Diestelhorst J, Beier R, Schäffer AA, Gertz EM, Schambach A, Kreipe HH, Pfeifer D, Engelhardt KR, Rezaei N, Grimbacher B, Lohrmann S, Sherkat R, Klein C. 2012. The phenotype of human STK4 deficiency. Blood 119:3450– 3457.

20. Nehme NT, Schmid JP, Debeurme F, André-Schmutz I, Lim A, Nitschke P, Rieux-Laucat F, Lutz P, Picard C, Mahlaoui N, Fischer A, de Saint Basile G. 2012. MST1 mutations in autosomal recessive primary immunodeficiency characterized by defective naive T-cell survival. Blood 119:3458–3468.

21. Sanchez-Vega F, Mina M, Armenia J, Chatila WK, Luna A, La KC, Dimitriadoy S, Liu DL, Kantheti HS, Saghafinia S, Chakravarty D, Daian F, Gao Q, Bailey MH, Liang W-W, Foltz SM, Shmulevich I, Ding L, Heins Z, Ochoa A, Gross B, Gao J, Zhang H, Kundra R, Kandoth C, Bahceci I, Dervishi L, Dogrusoz U, Zhou W, Shen H, Laird PW, Way GP, Greene CS, Liang H, Xiao Y, Wang C, Iavarone A, Berger AH, Bivona TG, Lazar AJ, Hammer GD, Giordano T, Kwong LN, McArthur G, Huang C, Tward AD, Frederick MJ, McCormick F, Meyerson M, Cancer Genome Atlas Research Network, Van Allen EM, Cherniack AD, Ciriello G, Sander C, Schultz N. 2018. Oncogenic Signaling Pathways in The Cancer Genome Atlas. Cell 173:321–337.e10.

22. Cagdas D, Halacli SO, Tan C, Esenboga S, Karaatmaca B, Cetinkaya PG, Balcı-Hayta B, Ayhan A, Uner A, Orhan D, Boztug K, Ozen S, Topaloglu R, Sanal O, Tezcan I. 2021. Diversity in Serine/Threonine Protein Kinase-4 Deficiency and Review of the Literature. J Allergy Clin Immunol Pract 9:3752–3766.e4.

23. Geng J, Sun X, Wang P, Zhang S, Wang X, Wu H, Hong L, Xie C, Li X, Zhao H, Liu Q, Jiang M, Chen Q, Zhang J, Li Y, Song S, Wang H-R, Zhou R, Johnson RL, Chien K-Y, Lin S-C, Han J, Avruch J, Chen L, Zhou D. 2015. Kinases Mst1 and Mst2 positively regulate phagocytic induction of reactive oxygen species and bactericidal activity. Nat Immunol 16:1142–1152.

24. Lee P-C, Machner MP. 2018. The Legionella Effector Kinase LegK7 Hijacks the Host Hippo Pathway to Promote Infection. Cell Host Microbe 24:429–438.e6.

25. Shehat MG, Aranjuez GF, Kim J, Jewett TJ. 2021. The Chlamydia trachomatis Tarp effector targets the Hippo pathway. Biochemical and Biophysical Research Communications 562:133–138.

26. García-Gil A, Galán-Enríquez CS, Pérez-López A, Nava P, Alpuche-Aranda C, Ortiz-Navarrete V. 2018. SopB activates the Akt-YAP pathway to promote Salmonella survival within B cells. Virulence 9:1390–1402.

27. Dacus D, Cotton C, McCallister TX, Wallace NA. 2020. Beta Human Papillomavirus 8E6 Attenuates LATS Phosphorylation after Failed Cytokinesis. Journal of Virology 94:10.1128/jvi.02184-19.

28. Wu SC, Grace M, Munger K. 2023. The HPV8 E6 protein targets the Hippo and Wnt signaling pathways as part of its arsenal to restrain keratinocyte differentiation. mBio 14:e01556–23.

29. Jr GG, Jeyachandran AV, Wang Y, Irudayam JI, Cario SC, Sen C, Li S, Li Y, Kumar A, Nielsen-Saines K, French SW, Shah PS, Morizono K, Gomperts BN, Deb A, Ramaiah A, Arumugaswami V. 2022. Hippo signaling pathway activation during SARS-CoV-2 infection contributes to host antiviral response. PLOS Biology 20:e3001851.

30. Wang Y, Xu X, Maglic D, Dill MT, Mojumdar K, Ng PK-S, Jeong KJ, Tsang YH, Moreno D, Bhavana VH, Peng X, Ge Z, Chen H, Li J, Chen Z, Zhang H, Han L, Du D, Creighton CJ, Mills GB, Cancer Genome Atlas Research Network, Camargo F, Liang H. 2018. Comprehensive Molecular Characterization of the Hippo Signaling Pathway in Cancer. Cell Rep 25:1304–1317.e5.

31. Yuan Z, Lehtinen MK, Merlo P, Villén J, Gygi S, Bonni A. 2009. Regulation of neuronal cell death by MST1-FOXO1 signaling. J Biol Chem 284:11285–11292.

32. Choi J, Oh S, Lee D, Oh HJ, Park JY, Lee SB, Lim D-S. 2009. Mst1-FoxO signaling protects Naïve T lymphocytes from cellular oxidative stress in mice. PLoS One 4:e8011.

33. Du X, Shi H, Li J, Dong Y, Liang J, Ye J, Kong S, Zhang S, Zhong T, Yuan Z, Xu T, Zhuang Y, Zheng B, Geng J-G, Tao W. 2014. Mst1/Mst2 regulate development and function of regulatory T cells through modulation of Foxo1/Foxo3 stability in autoimmune disease. J Immunol 192:1525–1535.

34. Zhou Y, Chen H, Liu L, Yu X, Sukhova GK, Yang M, Kyttaris VC, Stillman IE, Gelb B, Libby P, Tsokos GC, Shi G-P. 2017. Cathepsin K Deficiency Ameliorates Systemic Lupus Erythematosus-like Manifestations in Faslpr Mice. The Journal of Immunology 198:1846– 1854.

35. Hirai T, Kanda T, Sato K, Takaishi M, Nakajima K, Yamamoto M, Kamijima R, Digiovanni J, Sano S. 2013. Cathepsin K is involved in development of psoriasis-like skin lesions through TLR-dependent Th17 activation. J Immunol 190:4805–4811.

36. Adams JM, Cory S. 1998. The Bcl-2 Protein Family: Arbiters of Cell Survival. Science 281:1322–1326.

37. Kastan N, Gnedeva K, Alisch T, Petelski AA, Huggins DJ, Chiaravalli J, Aharanov A, Shakked A, Tzahor E, Nagiel A, Segil N, Hudspeth AJ. 2021. Small-molecule inhibition of Lats kinases may promote Yap-dependent proliferation in postmitotic mammalian tissues. 1. Nat Commun 12:3100.

38. Nishizawa M, Kataoka K, Goto N, Fujiwara KT, Kawai S. 1989. v-maf, a viral oncogene that encodes a “leucine zipper” motif. Proceedings of the National Academy of Sciences 86:7711–7715.

39. Soucie EL, Weng Z, Geirsdóttir L, Molawi K, Maurizio J, Fenouil R, Mossadegh-Keller N, Gimenez G, VanHille L, Beniazza M, Favret J, Berruyer C, Perrin P, Hacohen N, Andrau J- C, Ferrier P, Dubreuil P, Sidow A, Sieweke MH. 2016. Lineage-specific enhancers activate self-renewal genes in macrophages and embryonic stem cells. Science 351:aad5510.

40. Hegde SP, Zhao J, Ashmun RA, Shapiro LH. 1999. c-Maf Induces Monocytic Differentiation and Apoptosis in Bipotent Myeloid Progenitors. Blood 94:1578–1589.

41. Cao S, Liu J, Song L, Ma X. 2005. The Protooncogene c-Maf Is an Essential Transcription Factor for IL-10 Gene Expression in Macrophages1. The Journal of Immunology 174:3484–3492.

42. Xu Y, Xu M, Tong J, Tang X, Chen J, Chen X, Zhang Z, Cao B, Stewart AK, Moran MF, Wu D, Mao X. 2021. Targeting the Otub1/c-Maf axis for the treatment of multiple myeloma. Blood 137:1478–1490.

43. Graves JD, Draves KE, Gotoh Y, Krebs EG, Clark EA. 2001. Both Phosphorylation and Caspase-mediated Cleavage Contribute to Regulation of the Ste20-like Protein Kinase Mst1 during CD95/Fas-induced Apoptosis*. Journal of Biological Chemistry 276:14909–14915.

44. Palchaudhuri R, Lambrecht MJ, Botham RC, Partlow KC, van Ham TJ, Putt KS, Nguyen LT, Kim S-H, Peterson RT, Fan TM, Hergenrother PJ. 2015. A Small Molecule that Induces Intrinsic Pathway Apoptosis with Unparalleled Speed. Cell Reports 13:2027–2036.

45. Heimer S, Knoll G, Schulze-Osthoff K, Ehrenschwender M. 2019. Raptinal bypasses BAX, BAK, and BOK for mitochondrial outer membrane permeabilization and intrinsic apoptosis. Cell Death Dis 10:556.

46. Artemenko Y, Batsios P, Borleis J, Gagnon Z, Lee J, Rohlfs M, Sanséau D, Willard SS, Schleicher M, Devreotes PN. 2012. Tumor suppressor Hippo/MST1 kinase mediates chemotaxis by regulating spreading and adhesion. Proc Natl Acad Sci U S A 109:13632– 13637.

47. Hasselbring BM, Patel MK, Schell MA. 2011. Dictyostelium discoideum as a model system for identification of Burkholderia pseudomallei virulence factors. Infect Immun 79:2079– 2088.

48. Newton C, McHugh S, Widen R, Nakachi N, Klein T, Friedman H. 2000. Induction of Interleukin-4 (IL-4) by Legionella pneumophila Infection in BALB/c Mice and Regulation of Tumor Necrosis Factor Alpha, IL-6, and IL-1β. Infection and Immunity 68:5234–5240.

49. Wilkinson DS, Jariwala JS, Anderson E, Mitra K, Meisenhelder J, Chang JT, Ideker T, Hunter T, Nizet V, Dillin A, Hansen M. 2015. Phosphorylation of LC3 by the Hippo Kinases STK3/STK4 Is Essential for Autophagy. Molecular Cell 57:55–68.

50. Zheng Y, Wang W, Liu B, Deng H, Uster E, Pan D. 2015. Identification of Happyhour/MAP4K as Alternative Hpo/Mst-like Kinases in the Hippo Kinase Cascade. Dev Cell 34:642–655.

51. Meng Z, Moroishi T, Mottier-Pavie V, Plouffe SW, Hansen CG, Hong AW, Park HW, Mo J-S, Lu W, Lu S, Flores F, Yu F-X, Halder G, Guan K-L. 2015. MAP4K family kinases act in parallel to MST1/2 to activate LATS1/2 in the Hippo pathway. Nat Commun 6:8357.

52. Liu M, Tong Z, Ding C, Luo F, Wu S, Wu C, Albeituni S, He L, Hu X, Tieri D, Rouchka EC, Hamada M, Takahashi S, Gibb AA, Kloecker G, Zhang H, Bousamra M, Hill BG, Zhang X, Yan J. 2020. Transcription factor c-Maf is a checkpoint that programs macrophages in lung cancer. J Clin Invest 130:2081–2096.

53. Zaidi M, Troen B, Moonga BS, Abe E. 2001. Cathepsin K, osteoclastic resorption, and osteoporosis therapy. J Bone Miner Res 16:1747–1749.

54. van Schaarenburg RA, Magro-Checa C, Bakker JA, Teng YKO, Bajema IM, Huizinga TW, Steup-Beekman GM, Trouw LA. 2016. C1q Deficiency and Neuropsychiatric Systemic Lupus Erythematosus. Front Immunol 7:647.

55. Reid KBM. 2018. Complement Component C1q: Historical Perspective of a Functionally Versatile, and Structurally Unusual, Serum Protein. Frontiers in Immunology 9.

56. Ketelut-Carneiro N, Fitzgerald KA. 2022. Apoptosis, Pyroptosis, and Necroptosis—Oh My! The Many Ways a Cell Can Die. Journal of Molecular Biology 434:167378.

57. Creasy CL, Ambrose DM, Chernoff J. 1996. The Ste20-like Protein Kinase, Mst1, Dimerizes and Contains an Inhibitory Domain *. Journal of Biological Chemistry 271:21049–21053.

