## Supplementary Figures for "The Hippo kinases control inflammatory Hippo signaling and restrict bacterial infection in eukaryotic phagocytes"

A

| YAP1 Regulated Genes – Average Reads | | | | | |
|---|---|---|---|---|---|
| | WT | N5 | N12 | N13 | N19 |
| Ctgf | 0 | 0 | 0 | 0 | 0 |
| F3 | 0 | 0.4 | 0.2 | 0.8 | 0 |
| Cyr61 | 0.2 | 0.2 | 0.8 | 0.2 | 0 |
| Ankrd1 | 0.2 | 0 | 0 | 0.2 | 0 |
| Fjx1 | 0.2 | 0.2 | 0 | 0.2 | 1 |
| Foxf2 | 0.6 | 0.2 | 0 | 0.4 | 0.4 |
| Tgfb2 | 0.8 | 0.4 | 0.8 | 0.6 | 0 |
| Amotl2 | 1 | 1.2 | 0.6 | 2.2 | 1.2 |
| Igfbp3 | 1.2 | 1 | 1.2 | 1.8 | 0.8 |
| Ccdc80 | 1.6 | 1.6 | 1.4 | 4 | 1.4 |
| Nt5e | 2.4 | 2 | 3 | 2.2 | 0.6 |
| Crim1 | 6 | 6 | 6.8 | 7 | 6 |
| Gadd45a | 8.4 | 46.8 | 41 | 59.2 | 25.2 |
| Arhgef17 | 10.8 | 5.2 | 11.8 | 4.8 | 11.6 |
| Axl | 14 | 15.2 | 12.2 | 14.4 | 9.8 |
| Lats2 | 26.8 | 7.4 | 5.8 | 8 | 12 |
| Ptpn14 | 28.8 | 31 | 25.4 | 30 | 27.8 |
| Rbms3 | 142.4 | 143.6 | 103.8 | 159.4 | 114 |
| Nuak2 | 351.8 | 313.4 | 387.4 | 375 | 306.8 |
| Dock5 | 468.2 | 1002.2 | 900.4 | 1244 | 855 |
| Asap1 | 1051.2 | 686.8 | 664.2 | 860.4 | 577.6 |
| Myof | 1223.4 | 2942.6 | 2400.8 | 3706.2 | 3295.4 |

B

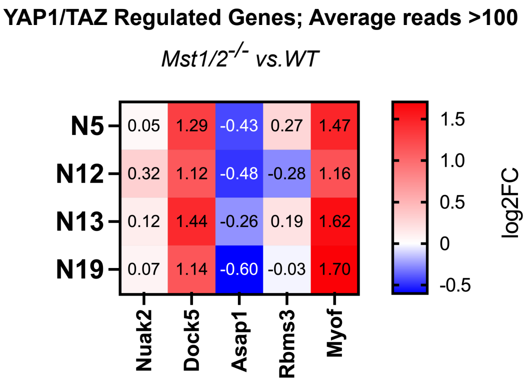


**Figure S1. Detection of YAP1/TAZ-regulated genes in macrophages by RNA sequencing.** **A.** Average transcript reads in iBMDMs by RNAseq for 22 established YAP1/TAZ-regulated genes based on Wang et al., 2018 (reference 30). The five genes with average reads >100 are highlighted in yellow boxes. **B.** Heat expression maps for the five YAP1/TAZ-regulated genes with average reads > 100 in $Mst1/2^{-/-}$ iBMDMs. The values within the boxes indicate the log2-fold changes of gene expression compared to WT iBMDMs.

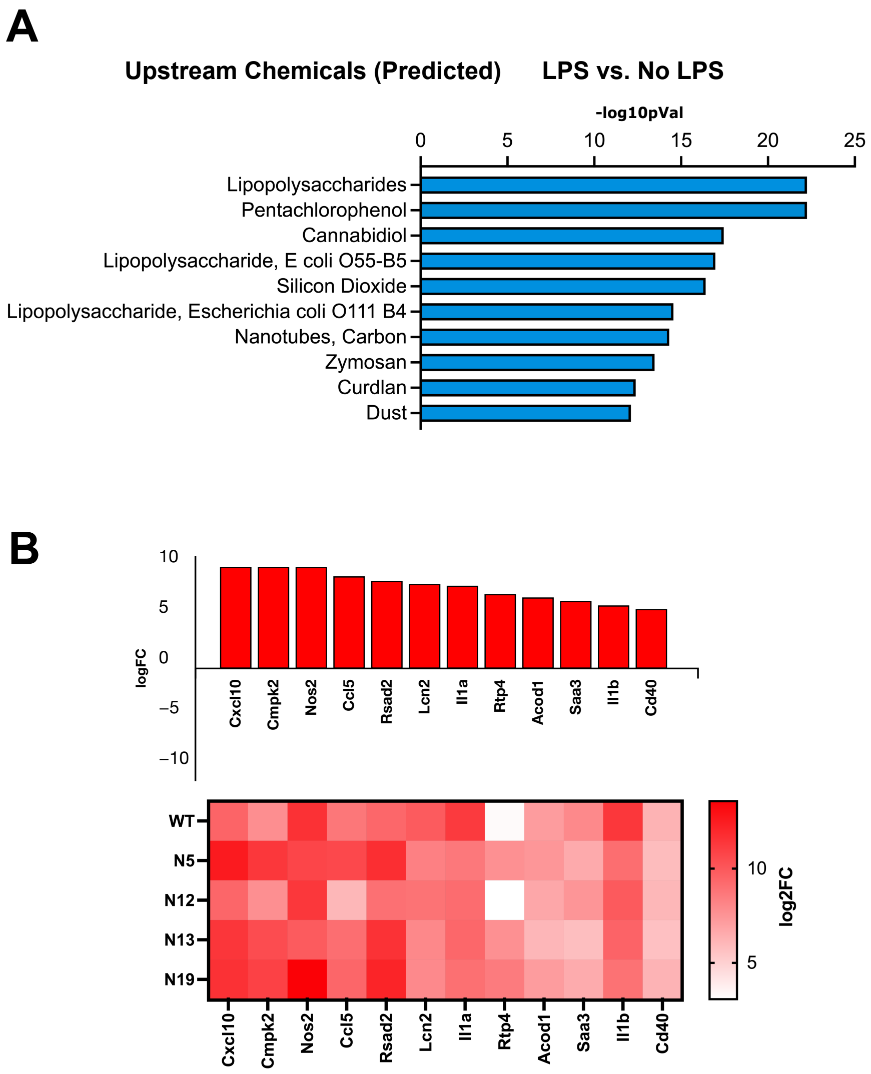


**Figure S2. WT and *Mst1/2*$^{-/-}$ iBMDMs respond to LPS treatment. A.** 706 differentially regulated genes by lipopolysaccharides (LPS) treatment in WT and *Mst1/2*$^{-/-}$ iBMDMs provided to iPathwayGuide. Upstream chemicals were predicted by iPathwayGuide and ranked by -log10 p value. This analysis confirms that iPathwayGuide accurately predicted LPS as the inducer for the differentially regulated genes in iBMDMs. **B.** Expression heat maps for the top 12 up-regulated genes in WT and *Mst1/2*$^{-/-}$ iBMDMs by LPS treatment. Similar levels of induction were observed in WT and *Mst1/2*$^{-/-}$ iBMDMs treated with LPS compared to no LPS treatment.

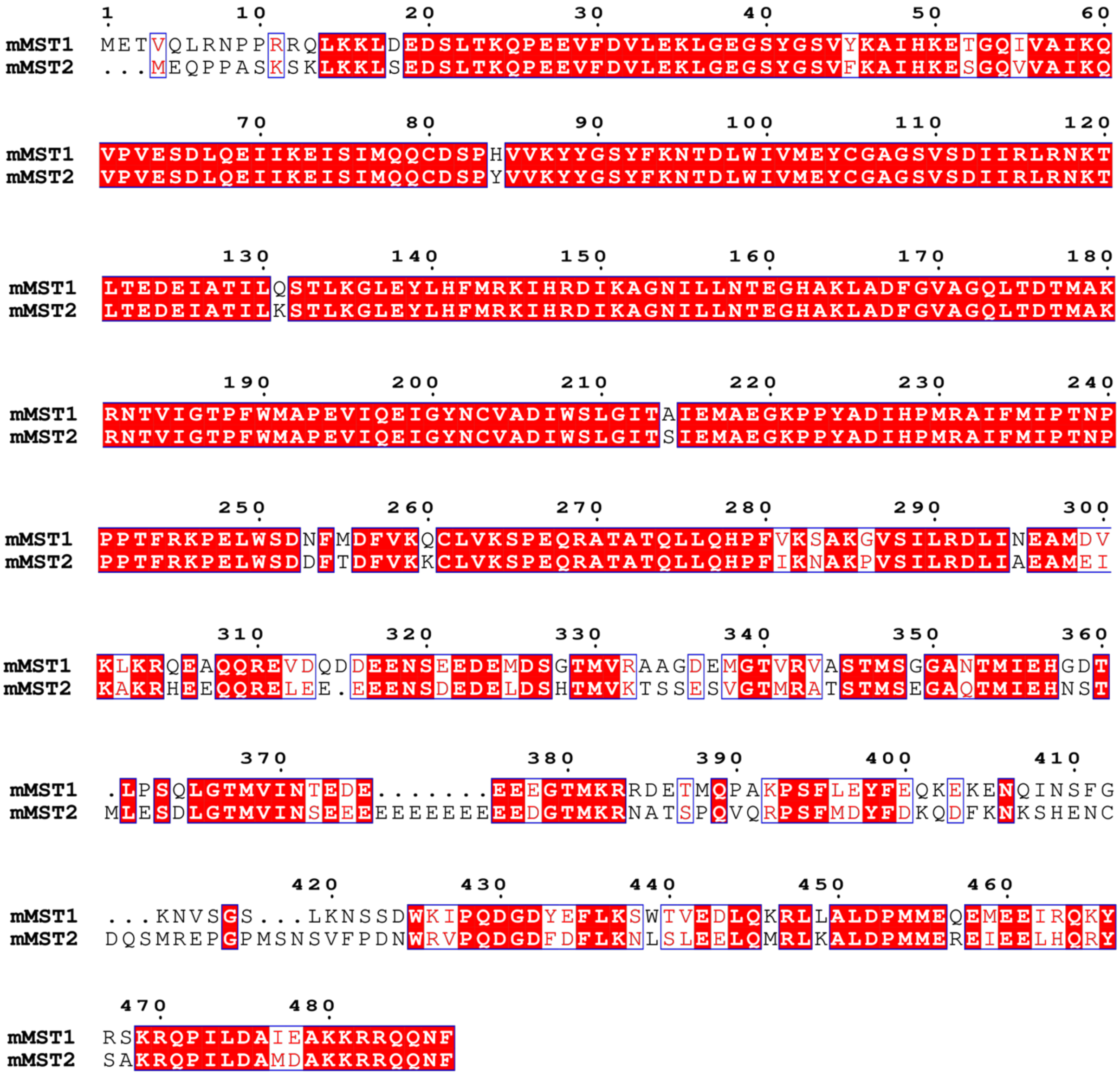

**Figure S3. Amino acid sequence alignments of MST1 and MST2. A.** Comparisons of amino acid sequences of mouse MST1 (NCBI Protein ID: NP_067395) and MST2 (NCBI Protein ID: NP_062609). Schematic diagrams generated by Clustal Omega (EMBL-EBI) and ESPript 3 (SBGrid). Identical amino acids are in the red background and white font. Similar amino acids are in the white background and red font. Numbers indicate the positions of amino acid residues of MST1.

**A**

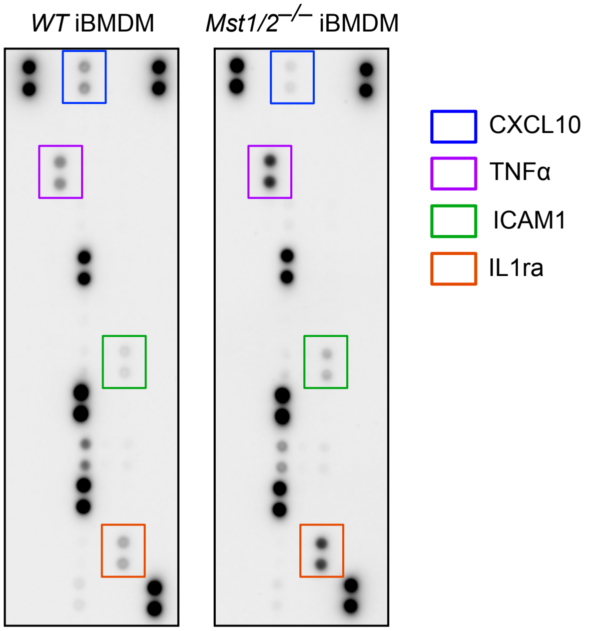


**B**

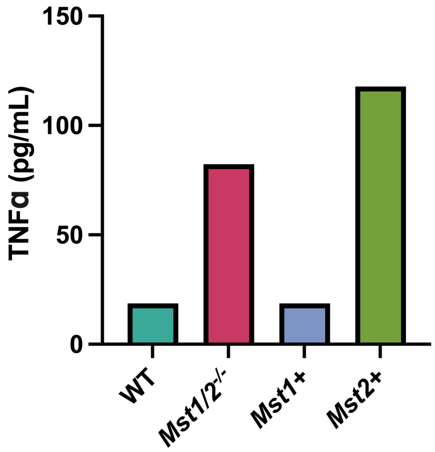


**Figure S4.** ***MST1/2 regulate secretion of cytokines into the conditioned media.* A.** Representative images of the cytokine arrays (R&D Systems) that detect 40 cytokines and chemokines in the conditioned media collected from WT and *Mst1/2*$^{-/-}$ iBMDMs challenged with *Lp02* for 3 hours. Array images shown are representative of three independent biological repeats. Cytokine or chemokine spots are manufactured in duplicate on the arrays, and the color boxes define the spots of cytokines or chemokines showing differential levels of secretion. **B.** WT, *Mst1/2*$^{-/-}$, *Mst1+*, and *Mst2+* iBMDMs were cultured for 24 hours and the conditioned media was collected. ELISAs were used to determine the concentration of TNFα in culture media between groups. Data shown is representative of four biological repeats.

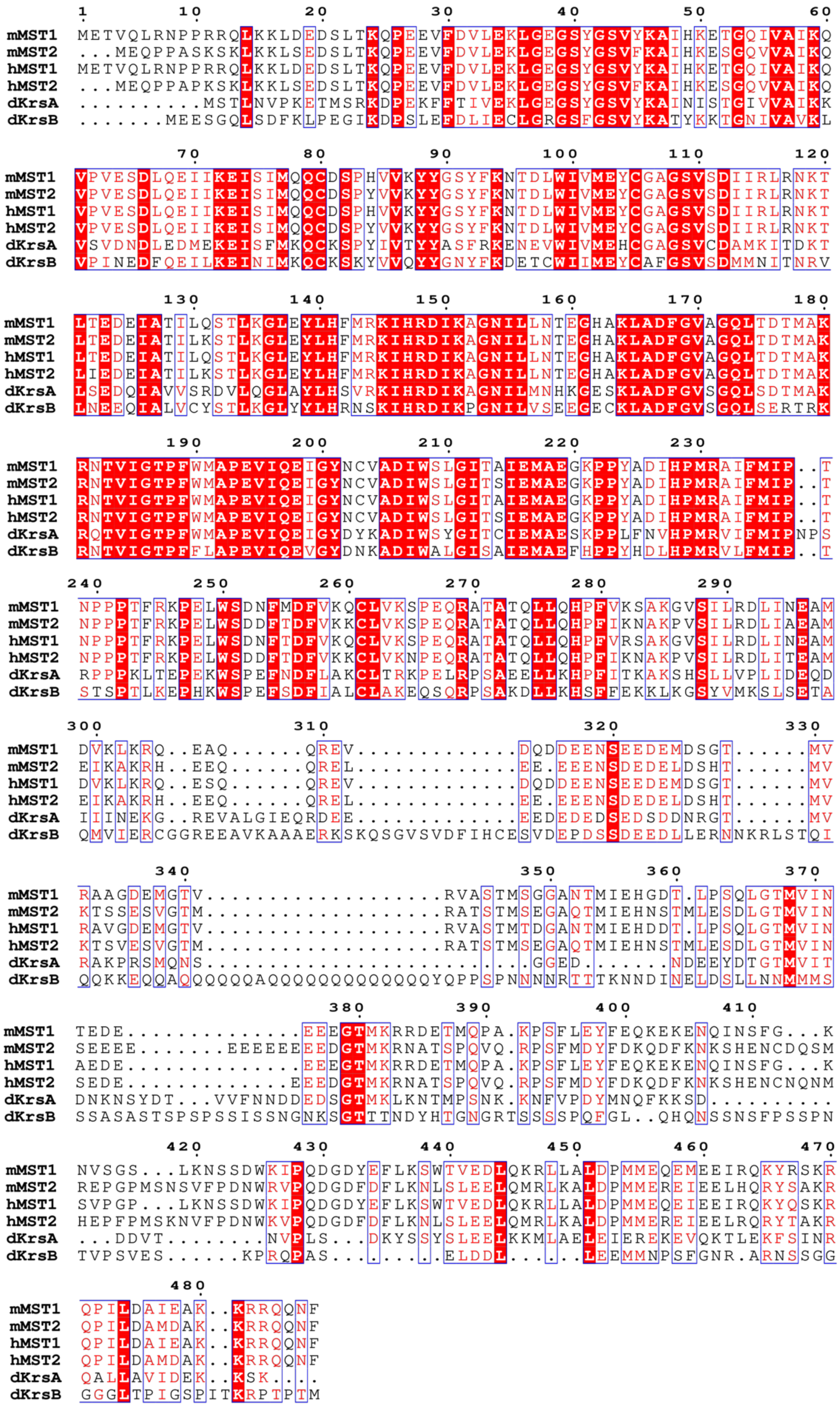
1 10 20 30 40 50 60
mMST1 METVQLRNPPRRQLKKLDEDSLTKQPEEVFDVLEKLGEGSYGSVYKAIHKETGQIVAIKQ
mMST2 ...MEQPPASKSKLKKLSEDSLTKQPEEVFDVLEKLGEGSYGSVFKAIHKESGQVVAIKQ
hMST1 METVQLRNPPRRQLKKLDEDSLTKQPEEVFDVLEKLGEGSYGSVYKAIHKETGQIVAIKQ
hMST2 ...MEQPPAPKSKLKKLSEDSLTKQPEEVFDVLEKLGEGSYGSVFKAIHKESGQVVAIKQ
dKrsA ..........MSTLNVPKETMSRKDPEKFFTIVEKLGEGSYGSVYKAINISTGIVVAIKK
dKrsB .......MEESGQLSDFKLPEGIKDPSLEFDLIECLGRGSFGSVYKATYKKTGNIVAVKL
70 80 90 100 110 120
mMST1 VPVESDLQEIIKEISIMQQCDSPHVVKYYGSYFKNTDLWIVMEYCGAGSVSDIIRLRNKT
mMST2 VPVESDLQEIIKEISIMQQCDSPYVVKYYGSYFKNTDLWIVMEYCGAGSVSDIIRLRNKT
hMST1 VPVESDLQEIIKEISIMQQCDSPHVVKYYGSYFKNTDLWIVMEYCGAGSVSDIIRLRNKT
hMST2 VPVESDLQEIIKEISIMQQCDSPYVVKYYGSYFKNTDLWIVMEYCGAGSVSDIIRLRNKT
dKrsA VSVDNDLEDMEKEISFMKQCKSPYIVTYYASFRKENEVWIVMEHCGAGSVCDAMKITDKT
dKrsB VPINEDFQEILKEINIMKQCKSKYVVQYYGNYFKDETCWIIMEYCAFGSVSDMMNITNRV
130 140 150 160 170 180
mMST1 LTEDEIATILQSTLKGLEYLHFMRKIHRDIKAGNILLNTEGHAKLADFGVAGQLTDTMAK
mMST2 LTEDEIATILKSTLKGLEYLHFMRKIHRDIKAGNILLNTEGHAKLADFGVAGQLTDTMAK
hMST1 LTEDEIATILQSTLKGLEYLHFMRKIHRDIKAGNILLNTEGHAKLADFGVAGQLTDTMAK
hMST2 LIEDEIATILKSTLKGLEYLHFMRKIHRDIKAGNILLNTEGHAKLADFGVAGQLTDTMAK
dKrsA LSEDQIAVVSRDVLQGLAYLHSVRKIHRDIKAGNILMNHKGESKLADFGVSGQLSDTMAK
dKrsB LNEEQIALVCYSTLKGLYYLHRNSKIHRDIKPGNILVSEEGECKLADFGVSGQLSERTRK
190 200 210 220 230
mMST1 RNTVIGTPFWMAPEVIQEIGYNCVADIWSLGITAIEMAEGKPPYADIHPMRAIFMIP..T
mMST2 RNTVIGTPFWMAPEVIQEIGYNCVADIWSLGITSIEMAEGKPPYADIHPMRAIFMIP..T
hMST1 RNTVIGTPFWMAPEVIQEIGYNCVADIWSLGITAIEMAEGKPPYADIHPMRAIFMIP..T
hMST2 RNTVIGTPFWMAPEVIQEIGYNCVADIWSLGITSIEMAEGKPPYADIHPMRAIFMIP..T
dKrsA RQTVIGTPFWMAPEVIQEIGYDYKADIWSYGITCIEMAESKPPLFNVHPMRVIFMIPNPS
dKrsB RNTVIGTPFFLAPEVIQEVGYDNKADIWALGISAIEMAEFHPPYHDLHPMRVLFMIP..T
240 250 260 270 280 290
mMST1 NPPPTFRKPELWSDNFMDFVKQCLVKSPEQRATATQLLQHPFVKSAKGVSILRDLINEAM
mMST2 NPPPTFRKPELWSDDFTDFVKKCLVKSPEQRATATQLLQHPFIKNAKPVSILRDLIAEAM
hMST1 NPPPTFRKPELWSDNFTDFVKQCLVKSPEQRATATQLLQHPFVRSAKGVSILRDLINEAM
hMST2 NPPPTFRKPELWSDDFTDFVKKCLVKNPEQRATATQLLQHPFIKNAKPVSILRDLITEAM
dKrsA RPPPKLTEPEKWSPEFNDFLAKCLTRKPELRPSAEELLKHPFITKAKSHSLLVPLIDEQD
dKrsB STSPTLKEPHKWSPEFSDFIALCLAKEQSQRPSAKDLLKHSFFEKKLKGSYVMKSLSETA
300 310 320 330
mMST1 DVKLKRQ..EAQ......QREV.............DQDDEENSEEDEMDSGT......MV
mMST2 EIKAKRH..EEQ......QREL.............EE.EEENSDEDELDSHT......MV
hMST1 DVKLKRQ..ESQ......QREV.............DQDDEENSEEDEMDSGT......MV
hMST2 EIKAKRH..EEQ......QREL.............EE.EEENSDEDELDSHT......MV
dKrsA IIINEKG..REVALGIEQRDEE............EEDEDEDSEDSDDNRGT......MV
dKrsB QMVIERCGGREEAVKAAAERKSKQSGVSVDFIHCESVDEPDSSDEEDLLERNNKRLSTQI
340 350 360 370
mMST1 RAAGDEMGTV...................RVASTMSGGANTMIEHGDT.LPSQLGTMVIN
mMST2 KTSSESVGTM...................RATSTMSEGAQTMIEHNSTMLESDLGTMVIN
hMST1 RAVGDEMGTV...................RVASTMTDGANTMIEHDDT.LPSQLGTMVIN
hMST2 KTSVESVGTM...................RATSTMSEGAQTMIEHNSTMLESDLGTMVIN
dKrsA RAKPRSMQNS........................GGED.......NDEEYDTGTMVIT
dKrsB QQKKEQQAQQQQQQAQQQQQQQQQQQQQYQPPSPNNNNRTTTKNNDINELDSLLNNMMMS
380 390 400 410
mMST1 TEDE...............EEEGTMKRRDETMQPA.KPSFLEYFEQKEKENQINSFG...K
mMST2 SEEEE.......EEEEEEEEDGTMKRNATSPQVQ.RPSFMDYFDKQDFKNKSHENCDQSM
hMST1 AEDE...............EEEGTMKRRDETMQPA.KPSFLEYFEQKEKENQINSFG...K
hMST2 SEDE..............EEEDGTMKRNATSPQVQ.RPSFMDYFDKQDFKNKSHENCNQNM
dKrsA DNKNSYDT...VVFNNDDEDSGTMKLKNTMPSNK.KNFVPDYMNQFKKSD..........
dKrsB SSASASTSPSPSSISSNGNKSGTTTNDYHTGNGRTSSSSPQFGL..QHQNSSNSFPSSPN
420 430 440 450 460 470
mMST1 NVSGS...LKNSSDWKIPQDGDYEFLKSWTVEDLQKRLLALDPMMEQEMEEIRQKYRSKR
mMST2 REPGPMSNSVFPDNWRVPQDGDFDFLKNLSLEELQMRLKALDPMMEREIEELHQRYSAKR
hMST1 SVPGP...LKNSSDWKIPQDGDYEFLKSWTVEDLQKRLLALDPMMEQEIEEIRQKYQSKR
hMST2 HEPFPMSKNVFPDNWKVPQDGDFDFLKNLSLEELQMRLKALDPMMEREIEELRQRYTAKR
dKrsA ...DDVT........NVPLS...DKYSSYSLEELKKMLAELEIEREKEVQKTLEKFSINR
dKrsB TVPSVES......KPRQPAS...........ELDDL.......LEEMMNPSFGNR.ARNSSGG
480
mMST1 QPILDAIEAK..KRRQQNF
mMST2 QPILDAMDAK..KRRQQNF
hMST1 QPILDAIEAK..KRRQQNF
hMST2 QPILDAMDAK..KRRQQNF
dKrsA QALLAVIDEK..KSK....
dKrsB GGGLTPIGSPITKRPTPTM

**Figure S5. Amino acid sequence alignments of the Hippo kinases in mouse, human and *D. discoideum* amoeba.** Comparisons of amino acid sequences among the Hippo kinases from *Mus musculus* (mMST1 NCBI Protein ID: NP_067395; mMST2 NCBI Protein ID: NP_062609), *Homo sapiens* (hMST1, NCBI Protein ID: NP_006273; hMST2, NCBI Protein ID: NP_006272) and *Dictyostelium discoideum* (dKrsA, NCBI Protein ID: XP_638650; dKrsB, NCBI Protein ID: XP_647461). Schematic diagrams generated by Clustal Omega (EMBL-EBI) and ESPript 3 (SBGrid). Identical amino acids are in the red background and white font. Similar amino acids are in the white background and red font. Numbers indicate the positions of amino acid residues of mMST1. We note that full-length KrsB has 1,105 amino acids and the N-terminal 526 residues with sequence homology to the other Hippo kinases are shown in the alignments.
